# Dynamics of Neutrophilia at the Neurovascular Unit Arising from Repeated Pulmonary Inflammation

**DOI:** 10.1101/2023.10.16.562508

**Authors:** Wesley Chiang, Herman Li, Linh Le, Jennifer David-Bercholz, Ana Caceres, Kamryn S Stecyk, Mariah Marrero, Amanda Pereira, Claire Lim, Danial Ahmad, James L McGrath, Ania K. Majewska, Niccolò Terrando, Harris A Gelbard

## Abstract

**Background:** The role of neutrophils in mediating neurovascular vulnerability has been increasingly implicated in various acute inflammatory models of neuroimmune crosstalk between the periphery and the brain. Whether neurovascular vulnerability is similarly modulated in the context of frequent, but not acute, inflammatory activation in the periphery is the aim of our study. Such a model of frequent inflammatory irritation is pertinent to understanding the neurologic risk of constant exposure to aerosolized environmental hazards leading to progressive pulmonary disease.

**Methods:** To model repeated pulmonary inflammation, we applied a three-dose regimen of intranasal (i.n.) lipopolysaccharide (LPS) in C57BL/6J mice and studied the impact on the inflammatory environment of the brain, with a specific focus on neutrophil dynamics at the neurovascular unit (NVU). Tissue and circulatory inflammatory profiles were screened via bronchoalveolar lavage (BAL) protein content and cellularity, transcript analysis of brain tissue, and flow cytometry of peripheral blood. Intravital two-photon microscopy (2PM) of the brain vasculature identified neutrophil dynamics at the NVU. Immunofluorescence validated neutrophil dynamics and identified neuroinflammatory hallmarks and peripheral immune factor interactions at the NVU. In vivo findings were corroborated and replicated in murine and human microphysiological systems (MPS) modeling the blood-brain barrier as a proxy demonstration of the translational relevance of our findings.

**Results:** 2PM of tdTomato-Ly6G+ neutrophils demonstrated increased levels of circulating neutrophils and corresponding engagement with the brain vasculature after the three-dose repeated i.n. exposure regimen. Neutrophilia at the NVU was corroborated with increased transcript levels of *Ly6G* and other pro-inflammatory markers. This coordination between endothelial physiology and neutrophil phenotypes was recapitulated in murine and human MPS models. System-wide neutrophilia in the lung and circulation was found to be cotemporaneous to neutrophilia at the NVU based on the cellularity of BAL and peripheral blood samples collected at the same endpoints. Immunohistochemical analysis of brain tissue implicates temporal coordination between vascular surface adhesion molecules with changes in neutrophil dynamics from adhesion, crawling, stalling, and transmigration. Extravasation of neutrophils was complemented by sustained paravascular deposition of fibrinogen and microgliosis up to 72 hours after the final i.n. dosing. Microglia-associated effector functions for synaptic pruning and regulation of neutrophil activity demonstrated distinct temporal profiles.

**Conclusions:** Our results identify systemic levels of neutrophilia accompanied by ingress and extravascular accumulation in brain parenchyma that correlated with sustained microglial activation. This neutrophil-centric lung-brain interaction is complemented by the observation of paravascular fibrinogen deposition that alters synaptic metabolism. Thus, we propose a mechanistic role for neutrophilia and associated inflammatory dysregulation as essential mediators along the lung-brain neuroimmune axis in a generalizable model of repeated respiratory exposure to inflammatory agents.

**Graphical Abstract:** 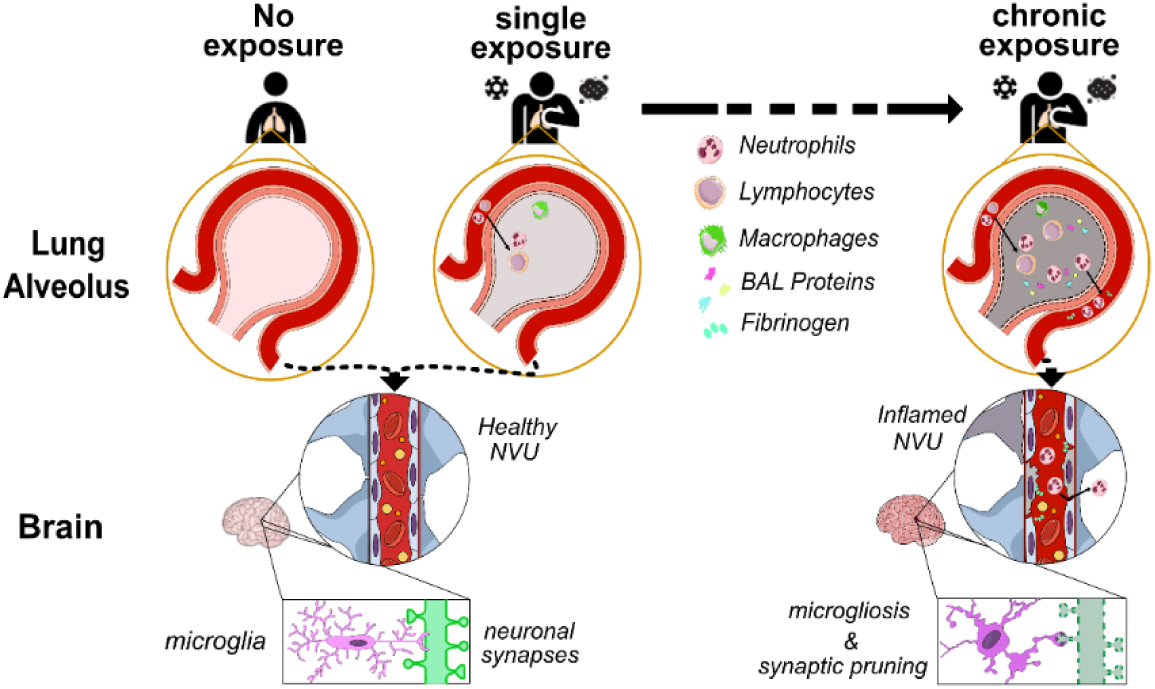

## Introduction

Respiration, an involuntary human function, is essential for our survival, yet also forces our bodies to be in constant exposure to potential health hazards (1). These hazards range from the increasing number of known and novel respiratory pathogens to also include aerosolized toxicants resulting from industrial byproducts (2). While both the skin and the respiratory system are constantly challenged by putative environmental hazards, direct access to the circulatory system is a vulnerability encountered by the pulmonary alveoli of the respiratory system but not the skin (3, 4). The surface area of the alveolar epithelium, roughly 25-times greater than the skin, serves as the primary barrier for safeguarding the lung and systemic circulation from potentially deleterious forms of inhaled inflammatory agents (5, 6). Under environmental conditions for either acute or chronic exposure to aerosolized inflammatory agents, respiratory inflammation may progress into severe cases known as acute lung injury (ALI) (2). The risk of hazardous environmental exposures in the air has only increased in densely populated, urbanized areas where individuals are continually exposed to a complex milieu of respiratory pathogens and aerosolized toxicants (3).

Fortunately, clinical presentation of respiratory symptoms for ALI are typically apparent and treated in an urgent fashion. However, the increased risk for subsequent complications to other organs, including the brain remains understudied. In fact, changes in neurologic function and brain pathology following ALI may not be immediate and are usually overshadowed by respiratory distress, and, as such, follow-up screening for cognitive dysfunction is not part of the standard of care (7, 8). A growing body of research in neuroimmunology has demonstrated considerable coregulatory mechanisms between the brain and peripheral organs, through which inflammatory activation at one end of the network likely indicates the presence of inflammation in the other (9). Given the expansive interface in the lungs for constant exposure to environmental toxicants, the respiratory system is likely an important site for host vulnerability that could implicate systemic dysregulation that ultimately alters the homeostatic functions in the brain (10).

The critical need to better understand the links between ALI and delayed onset of neurologic dysfunction was saliently highlighted by the recent COVID-19 pandemic; the plethora of “long COVID” syndromes, many of which include alterations in neurologic health, is only growing larger, with a report of 236,379 COVID-19 patients estimating that a 33-62% incidence of neuropsychiatric disease (11–15). The persistence of novel respiratory pathogens is likely to exacerbate the progressive decline in neurologic health associated with the pervasive exposure to environmental air pollutants (16). There is increasing evidence that continual exposure to aerosolized environmental toxicants biases vulnerable populations for enhanced susceptibility to neurodegenerative disorders, including Parkinson’s Disease and Alzheimer’s Disease, with acute delirium being another common complication that preludes subsequent neurodegeneration (4, 16–19). Coincidentally, a growing body of evidence points to the potential for respiratory pathogens to predispose individuals toward increased risk for onset of neurodegenerative diseases as well (8, 20).

Disturbingly, the full extent of neurological repercussion in subjects with sustained neuroinflammation after ALI will take years to emerge. These vulnerable groups range from individuals with postinfectious sequelae, such as “long COVID”, to those experiencing routine inflammation from environmental exposures (4, 14). Previous longitudinal studies between ALI and neurocognitive outcomes up to two years after hospital discharge have shown that the prevalence of neurocognitive impairment impacts nearly half the study population, with no improvement between years 1 and 2 (21). As these individuals age, their altered neuropathology due to ALI-associated neuroinflammation leaves them more susceptible to brain pathology, which further associates with worse prognosis, including mortality (22). This somber statistic highlights the importance of understanding early neuroimmune mechanisms that link ALI to brain inflammation.

To address this, we have developed an in vivo model of repeated respiratory inflammation via multi-hit intranasal (i.n.) delivery of lipopolysaccharide (LPS) to model cumulative induction of ALI relevant to densely populated urban areas. Here, we report a neutrophil-centric mechanism linking lung inflammation to disruption of neuroimmune homeostasis in the central nervous system (CNS). Despite being a bacterial endotoxin, LPS has been broadly validated as a generalized inflammatory agent for various disease models (23, 24). Via its canonical recognition by toll-like receptor 4 (TLR4), LPS signaling overlaps with the binding site of some viral envelope proteins, as well as many inflammatory signaling cascades encompassing both infectious and sterile injury through binding and downstream signaling at TLR4 sites (25–27). Additionally, LPS is commonly found as a component of biological organic matter in the complex composition of aerosolized particulate matter that are environmental toxicants to the respiratory system (28–31).

Our findings validate that multi-hit LPS model induces robust immune activation in the lung leading to systemic neutrophilia with neuroinflammatory activation in the cortex and hippocampus. Specifically, we identify neutrophil-mediated opening of the blood-brain barrier (BBB) accompanied by brain endothelial inflammation as a key instigator of damage to the neurovascular unit (NVU). These changes were mechanistically corroborated in murine-based microphysiological systems (MPS) modeling the contribution of neutrophils to BBB opening using murine endothelial cells and neutrophils isolated from the blood of C57BL/6J mice with ALI. In parallel, we validated and translated these observations with brain microvascular endothelial cells derived from human induced pluripotent stem cells (hiPSCs) cultured with freshly isolated human neutrophils in a human MPS model of the BBB. Lastly, we confirmed that neutrophilia and NVU damage were complemented by hallmarks of neuroinflammation, such as microgliosis and a decrease in postsynaptic elements, that involved direct interactions with systemic inflammatory hallmarks of neutrophilia, such as neutrophil crosstalk with microglia after ingress and deposition of paravascular fibrinogen. Thus, our results implicate a neutrophil-centric lung-to-brain neuroimmune axis linking ALI to subsequent brain inflammation.

## Results

### Establishing minimal repeat exposure paradigm for i.n. LPS induction of ALI-like pathology with coincident neuroinflammation

While chronic induction of severe lung damage, akin to ALI, has been extensively modeled in the literature (32–34), studies focused on the earliest phase of immune crosstalk along the lung-brain axis are limited (Figure 1A). Thus, we first determined the minimal number of i.n. exposures to subacute doses of LPS required to induce an ALI-like phenotype in C57BL/6J mice with a corresponding observation of inflammatory activation in the brain. 12-week-old mice were administered 40 μL i.n. LPS (0.25 μg/μL) every 24 hours, with mice randomly assigned to euthanasia and sample collection endpoints after 1, 2, and 3 doses of i.n. LPS (Figure 1B). Complementary saline instillation controls were conducted, and all cohorts of mice were euthanized 24 hours after the final dose of their assigned regimen for sample collection of either bronchoalveolar lavage (BAL) or whole brain tissue.

**Figure 1.**
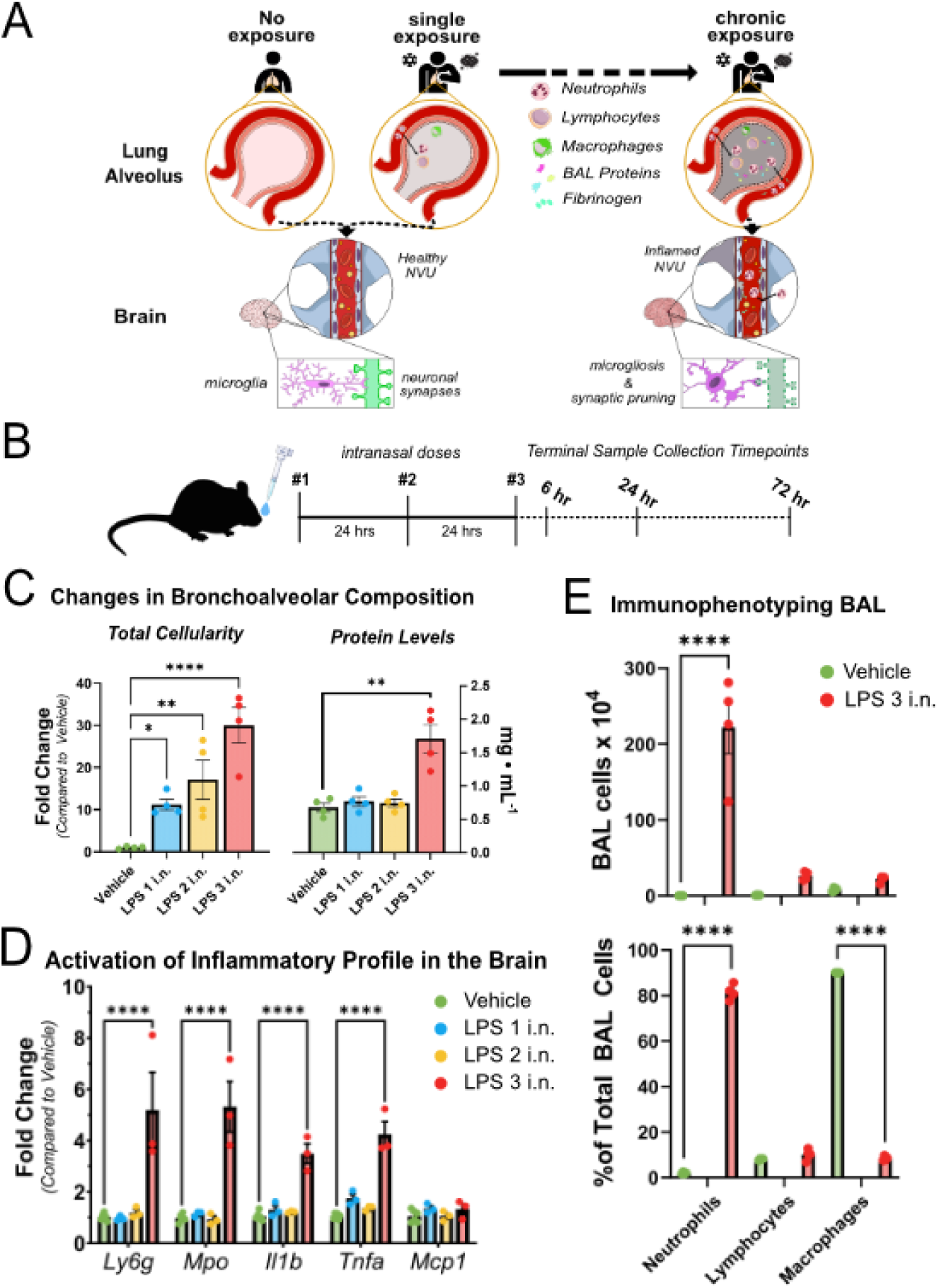
Multi-hit i.n. LPS drives robust ALI and sustains brain inflammation: **(A)** Repeat exposure paradigm for ALI pathology with ensuing systemic inflammatory profiles linking the lung to the brain. **(B)** Experimental design for three-dose regimen of i.n. administration paradigm of either saline vehicle or LPS. BAL and whole tissue sample collection occurred at a 24 hr after the final dose of assigned regimen. C57BL/6J male mice (12-weeks-old) were administered daily doses of 40 μL i.n. LPS (0.25 μg/μL in sterile saline) once (LPS 1 i.n.; 10 μg total), twice (LPS 2 i.n.; 20 μg total) or thrice (LPS 3 i.n; 30 μg total) with saline-matched controls. **(C)** Number of cells detected in BAL fluid was significantly increased following LPS 3 i.n. regimen, with robust increases in BAL total protein content. **(D)** Relative transcript abundance analyzed by qPCR of total RNA isolated from whole brain tissue homogenates. **(E)** Immunophenotyping via Diff-Quik stain to breakdown the specific composition of cell types that make up the BAL based on estimated raw totals and fractional composition of all cells counted. Data represented as mean ± SEM and analyzed with either one-way (total protein content) or two-way ANOVA of simple row effects (BAL cellularity and qPCR) and post-hoc correction with Holm-Sidak to determine statistically significant differences represented by asterisks for adjusted p-values that were found to be less than: 0.05(*), 0.01(**), 0.001(***), 0.0001(****).

The BAL was first screened for total cellularity and protein composition to evaluate the level of respiratory inflammatory following each i.n. LPS challenge (Figure 1C). Although all dosing regimens resulted in significant increases in total cellularity compared to matched saline controls (1 i.n. = 10-fold, 2 i.n. = 15-fold, 3 i.n. = 30-fold), an increase in total protein content in BAL was only observed after three exposures. The lack of significant changes in the proteinaceous content of the BAL after one and two doses of i.n. LPS suggest that the alveolar epithelium remains structurally intact and do not satisfy the criteria for induction of an ALI-like phenotype. Correspondingly, we evaluated the inflammatory profile of the brain by screening for changes in the transcript levels of a select panel of neuroinflammatory markers relevant to studies of chronic induction of ALI (Figure 1D). Similar to the profile of the BAL, only the three-dose regimen clearly demonstrated an inflammatory profile in the brain; we observed clear upregulation of transcripts for lymphocyte antigen 6 complex 6GD (*Ly6g*), myeloperoxidase (*Mpo*), interleukin 1β (*Il1β*), and tumor necrosis factor α (*Tnfα*). Of particular interest were the increases in *Ly6g* and *Mpo* in the brain given their primary association with neutrophil activation, an immune cell recruited during early phases of inflammatory activation. The lack of significant changes in the levels of *Mcp1* (monocyte chemotactic protein 1), typically released by activated microglia and tissue-resident macrophage, further confirms that the 3-dose regimen results in a linked lung-brain inflammatory profile during the early phases of ALI induced by repeat exposure to inflammatory agents.

### Cotemporaneous pulmonary, vascular, and cerebral neutrophilia in early-stage induction of ALI from repeat exposure model

We then phenotyped the specific cellular profile of the BAL in the LPS 3 i.n. paradigm to assess the inflammatory stage of the lung that is cotemporaneous with early stages of neuroinflammation (Figure 1E). Using a modified Wright-Giemsa stain (Diff-Quik), we identified neutrophils as the dominant constituent of the BAL (∼80% of all cells), and the primary cell type involved in driving the increased cellularity based on estimated raw counts (∼200-fold increase). Both lymphocytes and macrophages exhibited marginal, but not statistically significant, increases in estimated raw counts (∼12-fold to ∼7-fold, respectively), indicative of a neutrophil-dominant, early-phase inflammatory activation profile. The combined observation of neutrophil-dominant recruitment and a damaged, leaky lung epithelium (increased protein content in the BAL) after three exposures to i.n. LPS is consistent with early stages of severe ALI pathology (35, 36). The coincidence of early stages of inflammatory activation, indicated by neutrophilia, at both the lung and the brain suggest a neutrophil-mediated regulatory node that may shape the severity of neuroinflammation due to lung-related trauma.

To determine whether the cotemporaneous observations of neutrophilia at both the lung and the brain are linked, we screened if systemic neutrophilia was present in circulation to implicate lung-brain inflammatory crosstalk along a vascular route (Figure 2A). Peripheral blood samples collected at the same terminal endpoint (24 hours after 3^rd^ i.n. dose) were analyzed via flow cytometry and demonstrated that Ly6G+ cells are significantly increased compared to saline-matched controls, confirming our hypothesis of a vascular route for neutrophil-dependent lung-brain neuroimmune coupling (Figure 2B). Additionally, when we disaggregated our results into sex-specific samples, we observed evidence of neutrophilia in both male and female peripheral blood samples 24 hrs after the 3rd i.n. LPS dose; this indicates that systemic neutrophilia signaling in a lung-to-brain neuroimmune axis may regulate crosstalk in a sex-independent manner (37, 38).

**Figure 2:**
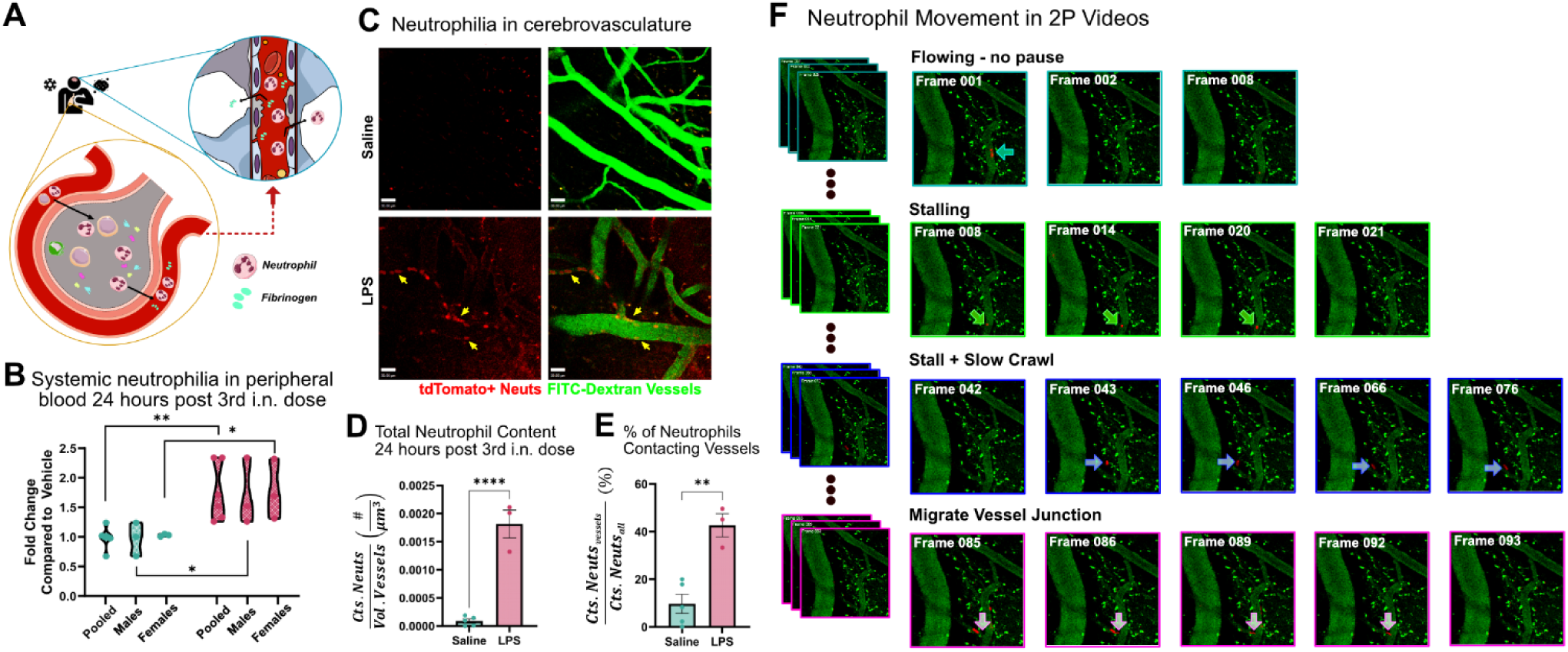
ALI induced by routine exposure paradigm drives systemic neutrophilia that reaches cerebral endothelium. **(A)** Schematic representation of systemic neutrophilia and fibrinogen in pulmonary circulation that reaches neurovascular units and subsequent transmigration or leakage in injured phenotypes. **(B)** Flow cytometry analysis of neutrophil content in peripheral blood of male and female C57BL/6J mice 24 hrs after the third i.n. dose of LPS compared to saline control. **(C)** Representative images of neutrophil engagement to brain endothelium obtained via 2P microscopy. Arrows represent neutrophil engagement to the vascular endothelium. Scale bar = 30 μm. The 2P time-course images were used to quantify **(D)** the average neutrophil density, defined as the total number of neutrophils normalized to vascular volume, and **(E)** the average fraction of total neutrophils that are contacting vessels. Values presented as mean ± SEM (saline n = 5; LPS n = 3). **(F)** Still frames taken from a representative 2P time-course of C57BL/6J mice treated with the 3x i.n. LPS ALI paradigm. Different sets of panels represent distinct functional features of neutrophil engagement with brain vasculature. Analysis for **(B)** was performed as a two-way ANOVA full-effect model with Holm-Sidak post-hoc correction. Analyses for **(D, E)** were performed as unpaired t-tests. Asterisks reflect significant comparisons where the adjusted p-value was found to be less than: 0.05(*), 0.01(**), 0.001(***), 0.0001(****).

We then translated the multi-hit i.n. LPS paradigm to C57BL/6-*Ly6g*(tm2621(Cre-tdTomato)Art e) (Catchup) reporter mice that express a fluorescent protein on one copy of their *Ly6g* loci to endogenously tag neutrophils on a similar genetic background to the mice used in our previous screens (39). The tdTomato-labeled neutrophils enabled the application of intravital two-photon (2P) microscopy through somatosensory cortical windows to characterize the specific dynamics of neutrophil behavior at the neurovasculature in real-time (Figure 2C). We intravenously (i.v.) injected 2,000 kDa FITC-dextran to outline the neurovasculature (green) and acquired 2P image sequences like Figure 2F, which demonstrated striking differences in tdTomato+ neutrophil (red) content within cerebral vessels of LPS 3 i.n. mice compared to the near-complete absence of neutrophilia in saline-matched controls. Quantitation of our 2P intravital data (Figures 2D, 2E) reveals that i.n. LPS induced robust increases in the total neutrophil count per µm^3^ in vessels (18-fold) and neutrophil contact with vessels (4-fold) when compared to i.n. saline. This quantification of increased neutrophil content in intracerebral vasculature corroborates the suggestion that systemic neutrophilia temporally links similar neutrophil profiles present in both the lung and brain during early stages of repeated pulmonary inflammatory activation.

### Multi-hit i.n. LPS induces neutrophil ingress and endothelial reactivity in the brain

The acquired 2P data allowed us to inquire about the biophysical behavior of neutrophils; the change in frequency of different types of neutrophil engagement with the endothelium could be indicative of distinct neutrophil activation states in response to the multi-hit i.n. LPS paradigm (40, 41). Via manual annotation of 2P image sequences from i.n. LPS exposed Catchup mice (Figure 2F), we frequently identified multiple states of neutrophil engagement at the endothelium – stalling, rolling/crawling, and transmigration through junctions – that we were unable to routinely identify in the vehicle (saline) videos (Supplemental Videos 1-4). Notably, neutrophil stalling that we observed in cerebral endothelium may reflect reduced blood flow in brain capillaries with adverse functional consequences (40) as well as sporadic neutrophil extravasation (41). This indication of distinct neutrophil activation states at the BBB, particularly transmigration through a vessel (Figure 3A), led us to harvest brains from the Catchup mice to specifically interrogate neutrophil behavior in brain regions we could not directly examine with intravital microscopy; i.e., in the vessels of the hippocampus. Neutrophil content (Ly6G+ DAPI+ objects, Figure 3B) in the stratum lacunosum moleculare (SLM) of the hippocampus were immunohistochemically quantified (Figure 3C) to provide support for our contention that neutrophilia in the BAL and circulation corroborated the corresponding neutrophilia we observed in cortical windows. Moreover, neutrophil ingress into the hippocampus further supports a hypothesis for neutrophil-mediated inflammatory linkage along a lung-vascular-brain axis during early stages of ALI, the severity of which may dictate the degree of decline in cognitive function that has been clinically documented in ALI patients (7, 8, 21, 22).

**Figure 3:**
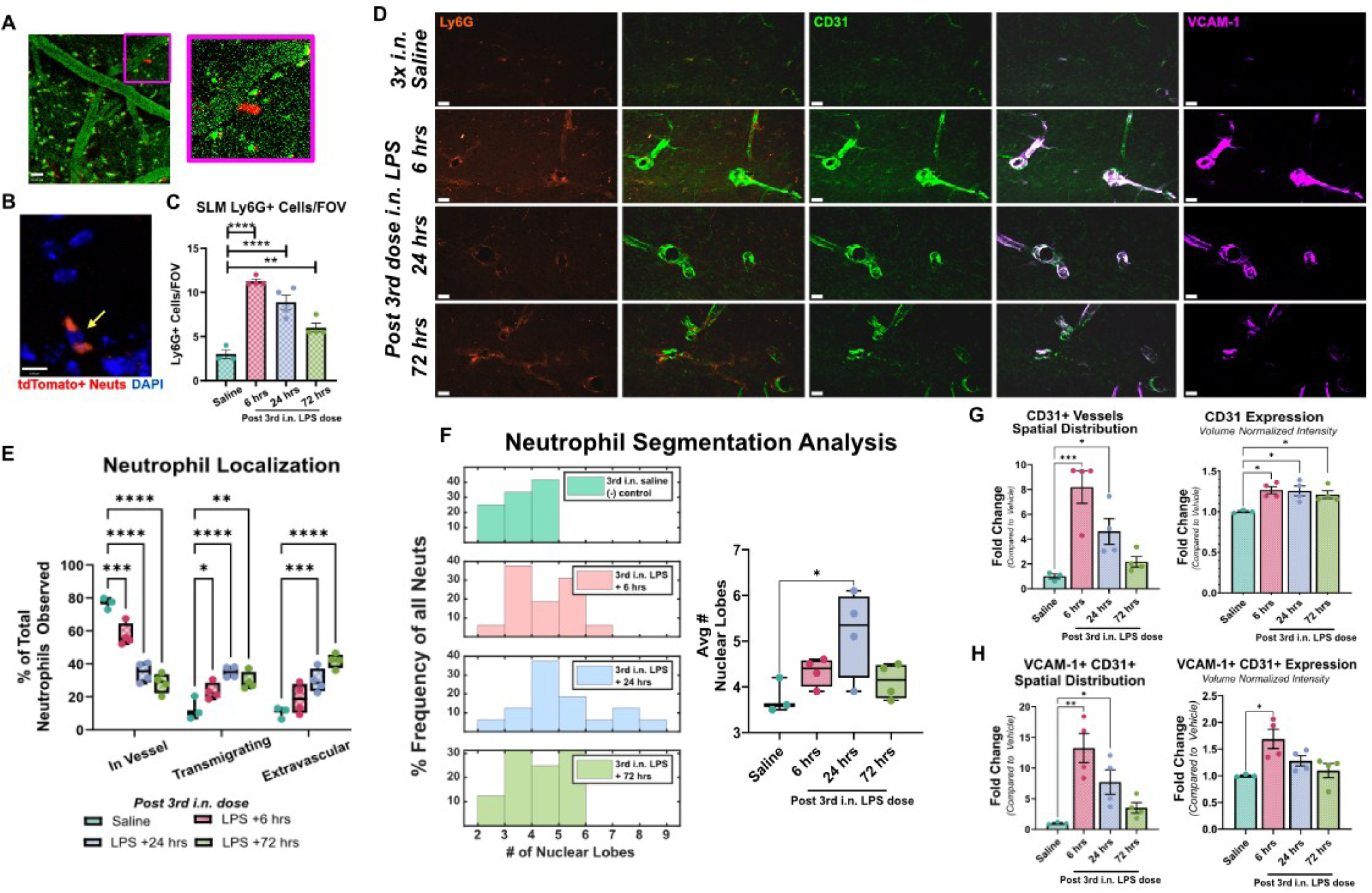
Enhanced engagement of neutrophils at neurovascular unit with inflamed vascular endothelium. **(A)** Still-frame image from 2P time-course demonstrating evidence of neutrophil extravasation out of brain vasculature into cortical parenchyma with digital magnification insert. Green (2000 kDa FITC-Dextran), Red (tdTomato-Ly6G). **(B)** Representative immunofluorescence of Ly6G+ cells in hippocampus of Ly6G-Catchup mice; red (tdTomato-Ly6G), blue (DAPI). Scale bar = 8 μm. **(C)** Ly6G+ cells/field of view (FOV) were quantified in the SLM of the hippocampus were quantified as mean ± SEM. **(D)** Representative immunohistochemical staining for tdTomato-Ly6G (orange), CD31 (green), and VCAM-1 (magenta) in Ly6G-Catchup mice. Scale bar = 20 μm. **(E)** Analysis of neutrophil localization from staining in **(D)** as to whether Ly6G+ DAPI+ neutrophil objects were in vessel, transmigrating, or extravascular based on co-incidence with CD31+ vessel objects. **(F)** Determination of nuclei segmentation in neutrophils at different time points post 3^rd^ i.n. dose of LPS represented as a histogram frequency distribution binned at integer number of nuclei per neutrophil, with summarized distribution. **(G, H)** Analysis of inflammation of brain vasculature indicated by distribution and expression levels of CD31 **(G)** as well as VCAM-1 in CD31+ vessel objects **(H)**. Analyses for **(C, F, G, H)** were performed using one-way ANOVA with Holm-Sidak post-hoc correction. Analysis for **(E)** was performed as a two-way ANOVA full effects model with Holm-Sidak post-hoc correction. All plots have all points plotted with either the mean ± SEM for bar plots and min-to-max for box-and-whisker plots (sham n = 3; LPS-treated groups n = 4). Asterisks reflect significant comparisons where the adjusted p-value was found to be less than: 0.05(*), 0.01(**), 0.001(***), 0.0001(****).

Next, we focused on hippocampal SLM, a relay between the entorhinal cortex and CA1 hippocampus, because of its involvement in neurodegenerative disease. In line with the known half-life of neutrophils, which can increase upon infiltration (42), we observed significant increases in neutrophils (tdTomato-Ly6G+/DAPI+ co-localized objects) at 6 hrs, 24 hrs, and 72 hrs after the 3^rd^ i.n. LPS dose compared to the three-dose i.n. saline control, with peak neutrophil counts observed at 6 hrs (43). Immunohistochemical analysis for a marker of endothelial cell interactions and immune cell trafficking, CD31 (also known as platelet endothelial cell adhesion molecule, PECAM) in the brain vasculature from the same postmortem tissue (Figure 3D) allowed further interrogation of spatial distribution of neutrophils at various terminal endpoints categorized as (i) tdTomato-Ly6G+ DAPI+ neutrophil objects in CD31+ vessels, (ii) transmigrating tdTomato-Ly6G+ DAPI+ neutrophil objects touching, but not in, CD31+ vessels, or (iii) extravascular tdTomato-Ly6G+ DAPI+ neutrophil objects not associated with CD31+ vessels (Figure 3E). This analysis demonstrated that while neutrophil counts peaked earlier (6 hrs) after the 3^rd^ i.n. LPS insult, the observation of neutrophil counts at later time points (24 hrs and 72 hrs) after the 3^rd^ i.n. dose can be attributed due to infiltrating neutrophils into the brain tissue itself, reflecting their increased half-life (42). Convincing evidence for this was reflected by the measured accumulation of extravascular neutrophil objects from 6 hrs to 72 hrs post 3^rd^ i.n. LPS dose.

We then assessed the presence of cellular indicators associated with neutrophil transmigratory behavior to corroborate our observations of increased ingress into the hippocampal parenchyma. Specifically, we focused on previous reports of neutrophil accumulation abluminal to the BBB associated with (i) increased nuclear lobe segmentation (44), (ii) available CD31 (or PECAM-1) adhesion sites (45), and (iii) VCAM-1 adhesion spots (45). Thus, expanding upon the analysis of our IHC staining in Figure 3D, we evaluated a semi-quantitative parameter for nuclear lobe segmentation in the neutrophils based on the number of DAPI+ objects touching Ly6G+ objects; this serves as a proxy for the number of nuclei associated with each neutrophil. Examining both aggregated frequency plots of total lobes per neutrophil and box plots of replicate averaged lobe numbers, we found that a significant increase in nuclear segmentation (i.e. “hypersegmentation”) was observed at the 24-hr terminal endpoint after the 3^rd^ i.n. LPS dose (Figure 3F). Additionally, concomitant increases in endothelial inflammation for CD31 expression remained elevated throughout the 72-hr period, but decreased in spatial distribution, potentially reflecting junctional “hot spots” in the BBB (Figure 3G). In contrast, VCAM-1 expression in CD31+ vessels reached a maximum at 6 hours after the 3^rd^ i.n. LPS exposure then declined to near baseline levels by the 72-hr endpoint. This decremental response may be due to negative transcriptional regulation of VCAM-1 expression by activating neutrophils or simple cleavage into soluble VCAM-1 that exacerbate endothelial damage (46, 47). Additionally, this downregulation in surface expression of VCAM-1 does not seem to be neither compensated by, nor dependent on, changes in GLUT-1 signaling (Supplemental Figure S1). In aggregate, these data are examples of widespread neutrophil engagement and sporadic migration through neurovascular endothelium in the somatosensory cortex that not only robustly manifests in a minimal repeat i.n. LPS exposure paradigm but is persistent beyond the typical early acute response period expected of neutrophils. These results imply a potential mechanistic role of cerebrovascular neutrophilia to both initiate and sustain the early phases of lung-brain inflammatory coupling.

### Neutrophil-induced endothelial dysfunction in murine and human microphysiological systems

To determine the translational relevance of our neutrophil-centric mechanism for brain inflammation via endothelial disruption, we used MPS with the μSIM microfluidic platform (48). A murine MPS (Figure 4) was first utilized to assess how well this model system recapitulates our observed in vivo phenotypes. After benchmarking the murine MPS, a human MPS (Figure 5) was used to assess functional parallels to the murine MPS, which, by proxy, serves to assess the relative likelihood that the neutrophil-centric neuroimmune mechanism linking cumulative induction of ALI to neuroinflammation may be extended to human disease.

**Figure 4:**
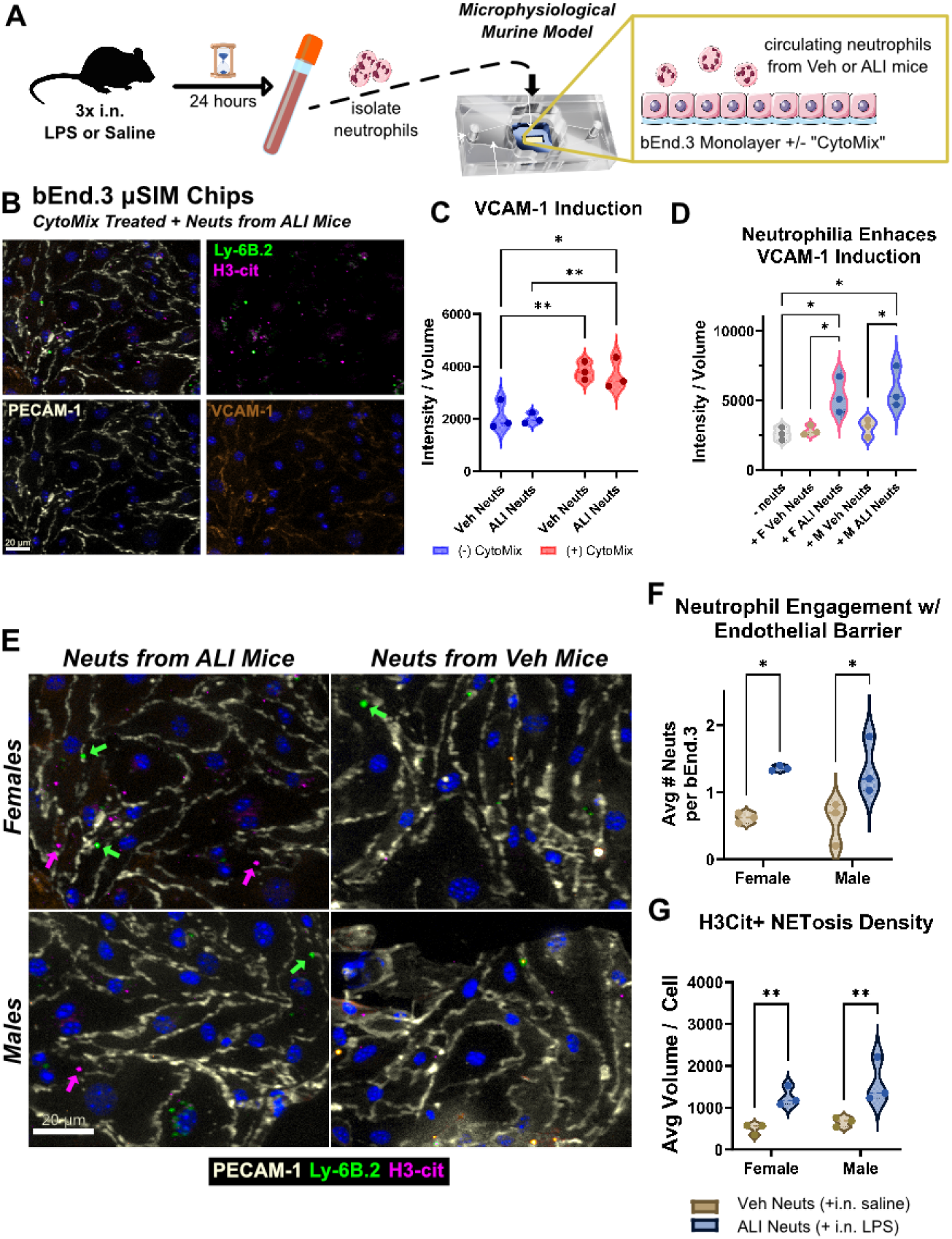
Recapitulation of neutrophil regulation of BBB health in repeat exposure inducted ALI-like paradigm in murine MPS. **(A)** Schematic diagram for isolation of saline vehicle or ALI neutrophil populations from mice given the same 3x i.n. paradigm from Figure 1B and introduced to a MPS for the murine BBB using cultured bEnd.3 monolayers stimulated with or without CytoMix. **(B)** Representative montage for detection of neutrophils and inflammatory activation of bEn3. Monolayers: White (PECAM-1), Orange (VCAM-1), Green (Ly6B.2), Magenta (H3-cit). Scale bar = 20 μm. **(C & D)** Inflammatory activation of bEnd.3 monolayers assessed by changes in VCAM-1 expression assessed by volume normalized intensity for VCAM-1+ objects. **(C)** Examines independent roles of CytoMix and inflammatory profile of neutrophils – from saline or LPS paradigm mice. **(D)** Determining whether sex-based differences exist from routine i.n. stimulation with or without LPS on CytoMix inflamed monolayers. **(E)** Representative montage for sex-based differences of neutrophil source in engaging with CytoMix stimulated murine MPS: White (PECAM-1), Green (Ly6B.2), Magenta (H3-cit). Scale bar = 20 μm. **(F)** Assessment of differential engagement of neutrophils due to sex as well as i.n. treatment paradigm. **(G)** Assessment of pro-inflammatory neutrophil phenotype via degranulation and NETosis from H3-cit+ objects. All plots show values for independent replicates (n = 3). Statistically significant differences were determined by Holm-Sidak post-hoc corrections after either one-way ANOVA (panel D) or two-way ANOVA fit with a full-effects model (panels C, F, G). These are represented by asterisks for adjusted p-values that were found to be less than: 0.05(*), 0.01(**), 0.001(***), 0.0001(****).

**Figure 5:**
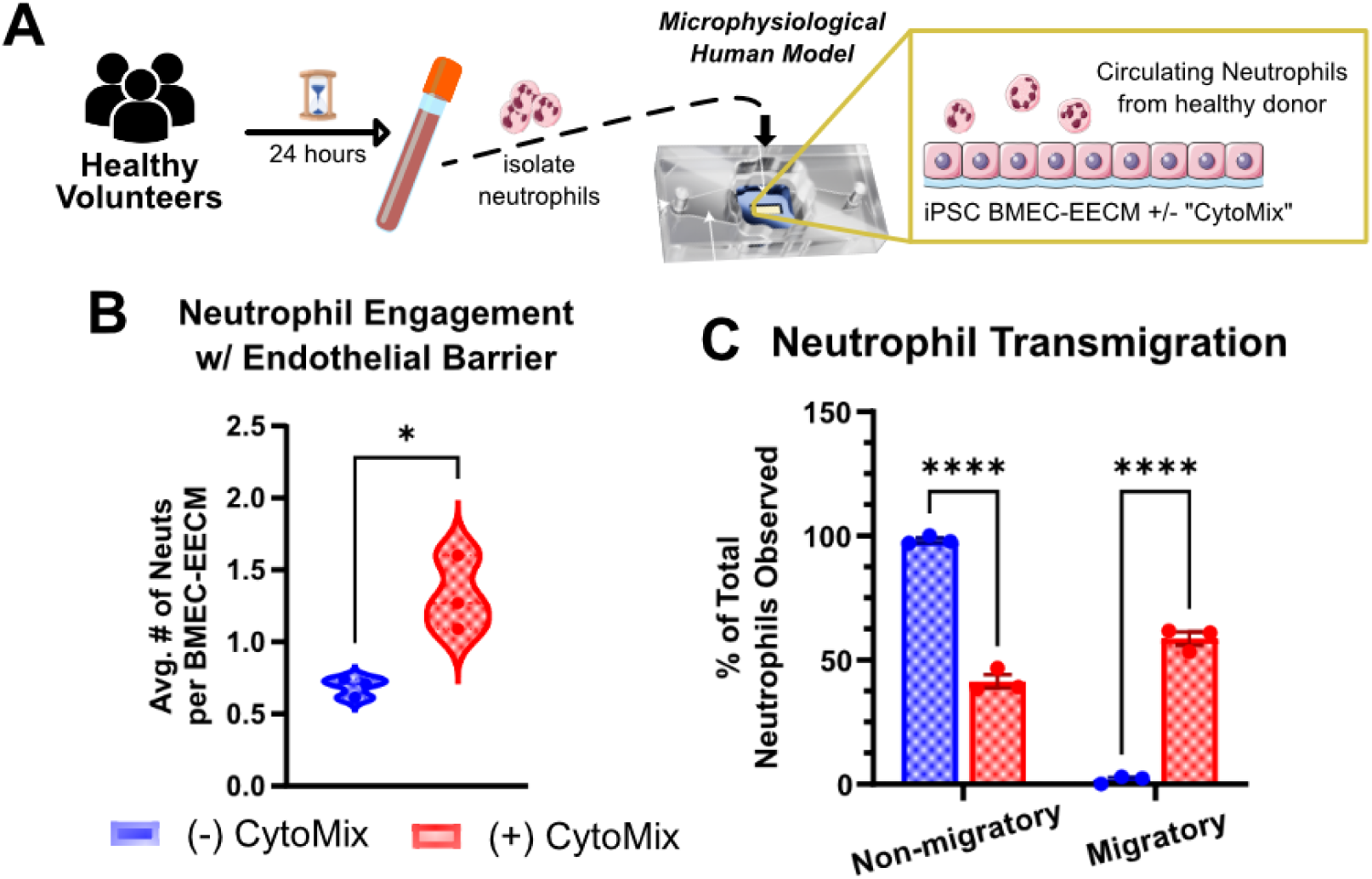
Translating neutrophil activity in repeat exposure ALI-paradigm to human-like model using MPS derived from hiPSCs. **(A)** A schematic overview of the human MPS constructed with EECM-BMECs derived from hiPSCs and exposed to neutrophils from human donors. **(B)** Analysis of neutrophil engagement at the EECM-BMEC monolayer in response to inflammatory activation and **(C)** resultant transmigration across the barrier.

In the murine MPS model (Figure 4A), bEnd.3 monolayers were cultured on optically transparent silicon membranes (µSiMs) and exposed to neutrophils isolated from the peripheral blood of C57BL/6J mice 24 hours after our three-dose repeated i.n. exposure paradigm with either vehicle (saline control) or LPS as outlined in Figure 1B. Neutrophils from saline or i.n. LPS groups reflect a vehicle baseline and stimulated inflammatory profile, respectively, and were applied to monolayer cultures of cerebral endothelial cells (bEnd.3) on a microphysiologic chip in the absence or presence of pro-inflammatory cytokines (“CytoMix”). The montage in Figure 4B depicts typical immunocytochemical results for expression of PECAM (CD31), VCAM-1 and citrullinated histone H3 (H3-Cit), a pivotal mediator of pro-inflammatory nuclear release of chromatin from neutrophils (neutrophil extracellular traps, NETs). We assessed the individual roles of a circulating inflammatory profile (i.e., systemic inflammation as a downstream effector of ALI) and neutrophil presence at the BBB in driving inflammatory activation, a predictor of BBB damage, in Figure 4C. In bEnd.3 models, basal expression of VCAM-1 is low, thus induction of increased expression of VCAM-1 is a standard proxy for inflammatory activation of the endothelial monolayer. Thus, VCAM-1 induction was assessed as the net increase of VCAM-1 expression in VCAM-1+ objects as volume-normalized intensity. This demonstrated that inflammatory induction of the endothelium was agnostic towards the activation profile of the neutrophils (i.e., isolated from vehicle vs ALI mice) without preliminary exposure to an inflammatory milieu, i.e., CytoMix. Furthermore, we investigated whether sex-based differences in neutrophil populations influenced our results from repeated i.n. exposure to saline or LPS; again, the sex of the neutrophil donors did not affect the agnostic behavior of the endothelium to the activation profile of the neutrophils (Figure 4D) without prior activation of the endothelial barrier.

After establishing that murine MPS could recapitulate endothelial activation in vivo, we then delineated neutrophil functionality at the barrier because of their pre-existing inflammatory profile. Neutrophil function was assessed as a measure of barrier engagement via Ly6B.2+ objects as well as identification of H3-cit+ objects (Figure 4E). Given that the VCAM-1 induction profiles demonstrated that the neutrophil functional profile does not alter barrier inflammation without prior stimulation, the neutrophil function studies were performed in MPS with only bEnd.3 monolayers pre-treated with CytoMix. In this paradigm, engagement with endothelial barriers was similar in both female and male mice (Figure 4F), with neutrophils isolated from ALI mice exhibiting increased engagement with the endothelium. Additionally, the density of H3-Cit+ objects, a proxy for NETosis density (Figure 4G), exhibited a similar trend.

We then translated our results into human MPS models using primary human neutrophils and hiPSC-derived endothelial cells (EECM-BMEC) grown in on the same µSiMs in the absence or presence of pro-inflammatory cytokines (Figure 5A). Like the murine MPS experiments, we delivered a pro-inflammatory cytokine milieu using CytoMix and reproducibly demonstrated differential neutrophil engagement with brain endothelial monolayers dependent on inflammatory induction (Figure 5B). After demonstrating similar physiological behavior between the murine and human MPS designs, we investigated differences in neutrophil transmigration across the µSiMs (Figure 5C). The results of this experiment confirm that pro-inflammatory induction of the EECM-BMEC monolayers by CytoMix increased migratory behavior of the neutrophils in agreement with the paradigm developed in our murine model system. In aggregate, the MPS data indicates that primed neutrophils in the vasculature augment inflammatory activation of the brain endothelium as a lung-brain neuroimmune linkage relevant to both murine and human systems.

### Paravascular fibrinogen accumulation and synaptic targeting accompany systemic neutrophilia during early lung-brain inflammatory coupling from repeat i.n. LPS exposure

We then returned to our mouse model to more extensively evaluate neutrophil-mediated changes at the NVU during early stages of lung-brain neuroimmune linkage. Particularly, the co-regulatory feedback between neutrophil activity at the endothelium with changes in fibrinogen metabolism is well documented in acute and chronic models of systemic inflammation with concomitant tissue-level inflammation (41, 49–54). In concert, these two inflammatory mediators are known to drive initial BBB breakdown, with paravascular fibrinogen eventually binding synapses with complement factors to act as biomolecular “eat me” signals for astrocytes and microglia (Figure 6A).

**Figure 6:**
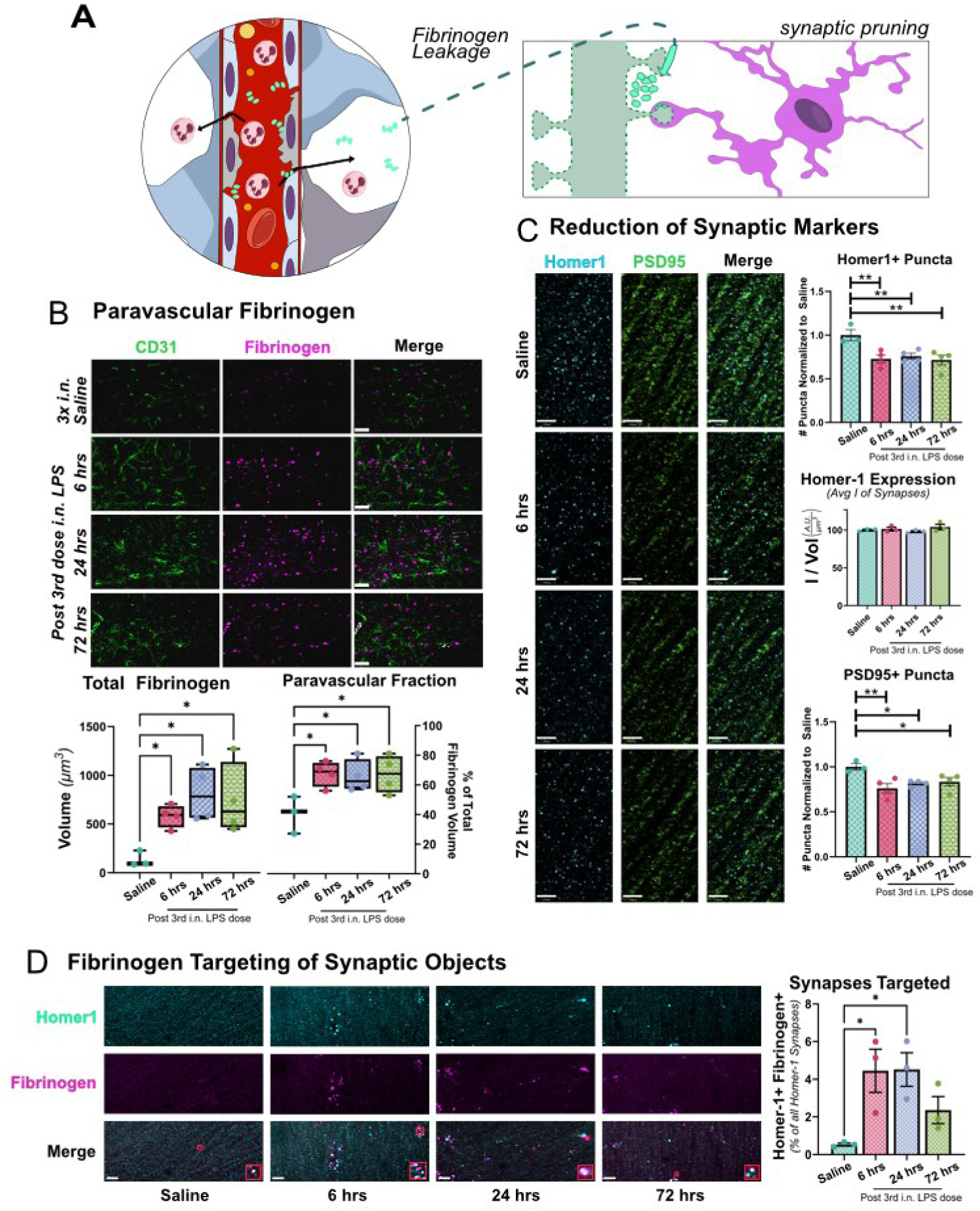
Systemic pro-inflammatory neutrophil profile for cumulative induction of ALI paradigm accompanied by fibrinogen-mediated damage to NVU. **(A)** Graphic representation of systemic neutrophilia and fibrinogenemia leading to leakage across NVU and damage to brain resident cells. **(B-D)** 12-week-old C57BL/6J mice received intranasal LPS described in Figure 1, followed by sacrifice 6, 24, and 72 hr after the last dose of LPS or saline. Brains were removed and processed for IF. (**B**) Montage of CD31 (green) and fibrinogen (magenta) expression in field-of-view, scale bar = 16 μm. Total and paravascular fibrinogen levels as a function of time after 3^rd^ i.n. LPS exposure **(C)** Montage of pre-synaptic Homer-1 (cyan) and post-synaptic PSD-95 (green) objects, scale bar = 20 μm. Changes in total synaptic architecture measured at 6 hrs, 24 hrs, and 72 hrs after final i.n. exposure. **(D)** Analysis of direct interaction between fibrinogen (magenta) and synapses via Homer-1 (cyan), scale bar = 15 μm. All quantifications have all data points plotted (saline n=3; LPS-treated groups n=4) and were performed as one-way ANOVA with Holm-Sidak post-hoc analysis to determine statistically significant differences represented by asterisks for adjusted p-values that were found to be less than: 0.05(*), 0.01(**), 0.001(***), 0.0001(****).

Upon staining for fibrinogen at the hippocampus, we found demonstrable increases in deposition, assessed as the total volume of immunoreactive objects, around the vasculature that is persistent throughout the 72-hr observation window after the third i.n. LPS exposure. This increase of fibrinogen+ objects was complemented by a similarly sustained decrease in spatial overlap with CD31 expression, reflecting paravascular deposition in the parenchyma (Figure 6B). We corroborated this assessment of BBB leakage via analysis of low molecular weight dextran extravasation in the hippocampus and identified a significant increase in leakage from big vessels (Supplemental Figure S2A). We also measured changes in low molecular weight dextran extravasation in the OB as a potential byproduct of i.n. stimulation but found no significant differences in leakage from neither small nor large vessels at the terminal endpoints involved in this study (Supplemental Figure S2B).

Paravascular deposition of fibrinogen is typically correlated with targeting of synapse with complement factors as biomolecular “eat me” signals for microglia and astrocytes to engulf and eliminate (Figure 6A) that leads to deleterious alterations in cognitive function (51, 55). Investigation of synaptic structures labeled by post-synaptic density 95 (PSD95) and scaffolding protein Homer-1 in the HPC (Figures 6C) identified consistent downregulation in the number of detected puncta for both markers compared to saline vehicle. Examination of the average expression of Homer-1 per puncta, by normalizing the intensity of each punctum to its volume, demonstrated no significant differences, and represents an “all-or-nothing” loss of post-synaptic features. Linking this to fibrinogen deposition and synaptic elimination (55), we find significant colocalization of fibrinogen immunoreactive objects with Homer-1 immunoreactive puncta suggesting that the increases in paravascular fibrinogen are associated with loss of postsynaptic elements (Figure 6D).

### Persistent Neutrophil ingress correlates with detuning of microglial activations profiles

The decline in synaptic densities coincident with increased paravascular deposition of fibrinogen implies the activation of glial cells, a typical feature of neuroinflammatory activation (41, 51, 52, 56). Interestingly, when evaluated for IBA1 and CD68 immunoreactivity as proxies for inflammation and phagocytic activity in microglia (Figure 7A), we found a temporal profile that demonstrated peak microglial activation between 6 and 24 hours after the third i.n. LPS dose, followed by a notable decline in both IBA1+ and CD68+ objects at the 72-hour endpoint (Figures 7B and 7C). This temporal profile was captured at both an ensemble aggregate level, as well as population-level distribution of individually segmented IBA1+ microglia cells (Supplemental Figure S3). The timing of this decline in microgliosis and phagocytic activity, however, may explain the sharp loss of synaptic content within 6 hrs after the third i.n. LPS, that then stagnates through the 72-hr endpoint (Figure 6C). Given the similar levels of fibrinogen-targeted Homer-1 densities (Figure 6D), a continued decline in synaptic densities was expected through the 24-hr timepoint; this suggests a potential detuning of phagocytic potential in the microglia.

**Figure 7:**
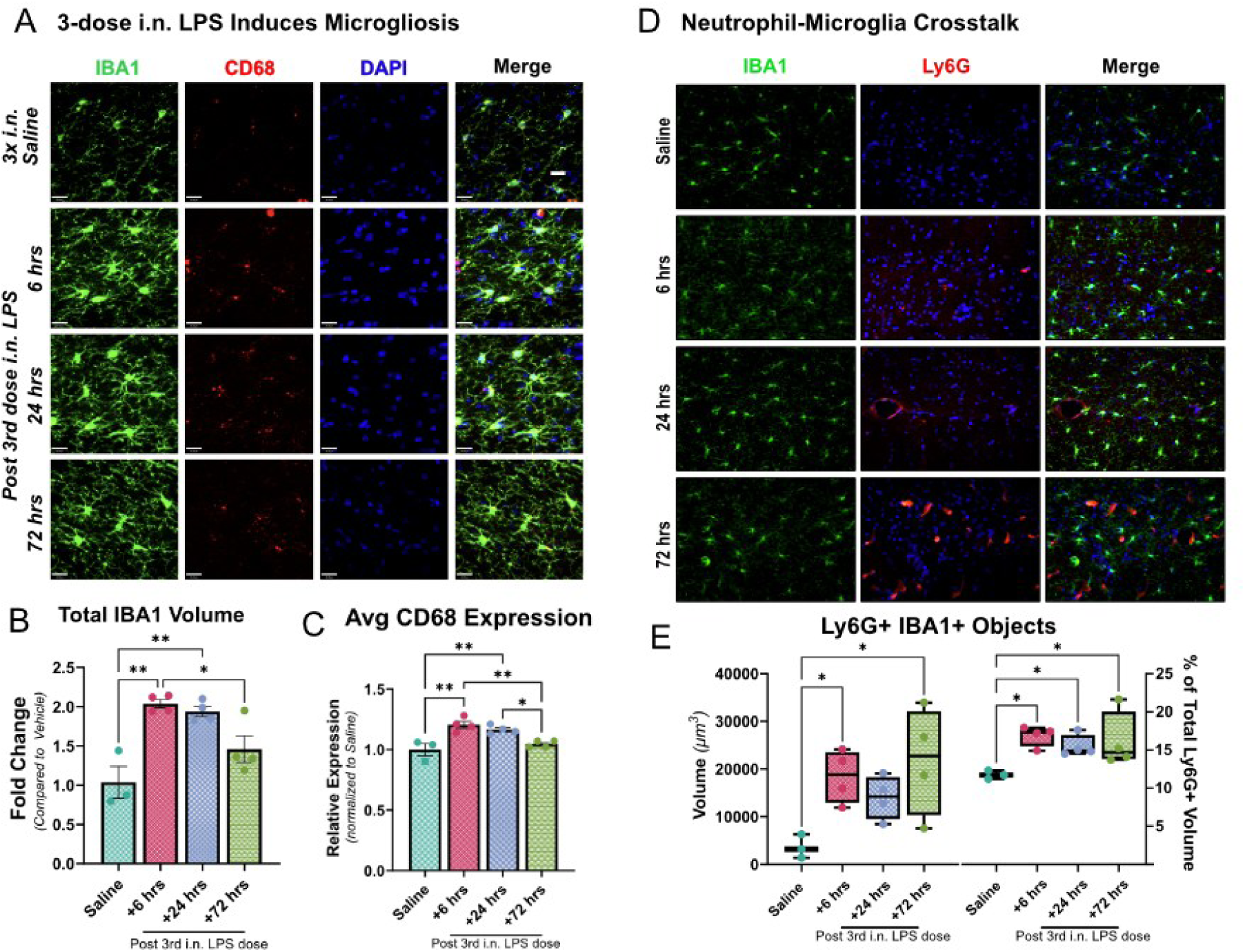
Microglial interactions with Neutrophils. 12-week-old Catchup mice received either a three-dose regimen of either i.n LPS or saline, followed by terminal sample collection at 6 hrs, 24 hrs, and 72 hrs after the final dose of the assigned regimen. Whole brains were harvested and processed for IF. (A) Microglial montage: Green (Iba1), red (CD68), blue (DAPI). (B) Iba1 intensity normalized to saline control over field-of-view. (C) CD68 total intensity normalized to Iba1 total volume. over field-of-view. Values presented as mean ± SEM (saline n=3; LPS-treated groups n=4). (D) Neutrophil-microglia interaction montage: Green (IBA1), red (Ly6G), blue (DAPI). (E) Analysis of neutrophil-microglia interactions by screening for segmented objects that are double positive for Ly6G and IBA1, represented as total volume of such objects and relative fraction of all Ly6G+ objects. Each data point reflects the average of a biological replicate’s hippocampus sampled over two regions in the SLM on each hemisphere over two to three different sections of the same brain. Analyses were performed as one-way ANOVA with Holm-Sidak post-hoc correction to determine statistically significant differences represented by asterisks for adjusted p-values that were found to be less than: 0.05(*), 0.01(**), 0.001(***), 0.0001(****).

This decline in microglial activation could simply arise from transition to an inflammatory resolving state, such as efferocytosis of apoptotic cells and their associated debris (i.e., neutrophils), after a peak inflammatory stage (57–60). This may exhaust the phagocytic surveillance of microglia away from fibrinogen-targeted synapse. However, the decline in CD68 immunoreactivity does not align with the phagocytic clearing of debris in efferocytosis (Figure 7C). Furthermore, the kinetics of neutrophilia up to 72 hours after the third i.n. LPS exposure suggests continued ingress through the BBB (Figure 3) that upon interaction with microglia could either drive sustained pro-inflammatory activation in the parenchyma (61, 62) or drive immunosuppression of certain microglial activation states relevant to cognitive impairment (63, 64). In both scenarios, however, neutrophil engagement with microglia has been shown to downregulate phagocytic function. Thus, we screened for whether the persistent accumulation of neutrophils in the parenchyma resulted in significant levels of neutrophil-microglia crosstalk that may explain the temporal profile of microglial function and synaptic loss in our previous data. As shown in Figures 7D and 7E, cellular objects that contain regions positive for both Ly6G and IBA1 fluorescence increases at the +6 hrs endpoint and is persistent through the +72 hrs endpoint after the third i.n. LPS exposure. The kinetics of this may explain the rapid loss of synapses that plateaus after +6 hrs, with a corresponding decline in microglial phagocytic capacity at +24 hrs.

## Discussion

Frequent, to almost constant, exposure to aerosolized inflammatory agents, such as seasonal respiratory pathogens and particulate matter in air pollution can lead to severe respiratory inflammation, a chronic corollary to ALI with consequential changes to brain inflammatory health (3). Despite the clear evidence for lung-brain neuroimmune coupling during severe pathology in chronic models, the exact mechanism that initiates such linkage between the two tissues is missing. As such, we used a subacute respiratory exposure of LPS to identify the minimal number repeated exposures required to both induce a severe lung injury phenotype with coincident changes in brain inflammatory profile that is bilateral and seemingly sex-independent (Supplemental Figure S4). LPS was chosen as a generalizable stimulus for ALI due to how it overlaps in inflammatory signaling cascades with other microbial virulence factors and some viral envelope proteins; it is also commonly found as one of the bio-organic matter in the complex, heterogeneous mixture of components in nano-/micro-particulate matter involved in pro-inflammatory stimulation of the respiratory tract (28–31). Using this paradigm, we investigated the early stages of lung-to-brain inflammatory coupling via a vascular route mediated by cotemporaneous neutrophilia observed in the lung, circulation, and brain.

By evaluating changes in BALF cellularity and proteinaceous content, we demonstrated that lung barrier breakdown, indicative of severe lung injury, occurred after three daily i.n. exposures of LPS. Interestingly, a measurable difference in inflammatory transcripts from brain tissue was also apparent after the same exposure time course. Further analysis of the BALF cellular composition identified extensive neutrophilia in the lungs, with corresponding increases in neuroinflammatory transcripts associated with neutrophil function (Ly6G, MPO), without significant changes in transcripts for monocyte chemoattractant protein 1 (MCP-1) despite its prevalence as a general chemoattractant for immune cells (65). This implied a chemoattractant milieu that is likely neutrophil-specific, in-line with our goal of defining a minimal subacute i.n. LPS exposure paradigm to investigate early-stages of lung-brain neuroimmune coupling relevant to understanding the effects extended chronic exposure.

The observation of neutrophilia in both the lung and the brain led us to investigate a neutrophil-centric mechanism characterized by systemic contributions to neuroimmune dysregulation resulting in damage to the NVU with implications for altered cognitive function. This supported a potential substrate of “brain fog” and cognitive dysfunction commonly reported after ALI, as well as the purported links of inhaled inflammatory agents with increased susceptibility for dementia in vulnerable populations living in dense or industrialized urban regions (3, 4, 8, 16, 17, 20). Via a combination of flow cytometry, 2P intravital microscopy, and IHC of thin brain slices, we reproducibly measured increased neutrophil content in peripheral circulation and at the NVU to confirm a systemic neutrophilia profile along a vascular route to link the lung and brain inflammatory states during early stages repeated subacute respiratory exposure. Annotating neutrophil behavior in the collected 2P image sequences allowed us to document instances of different neutrophil-endothelial cell interactions such as stalling, rolling, and even transmigration across a junction. Complementary IHC analysis of brain tissue to quantify the fractional content of neutrophils that are in vessels, transmigrating, or present in extravascular brain parenchyma further supported our observation of increasing extravascular neutrophils over a 72-hr window post 3^rd^ i.n. dose of LPS and is congruent with documented neutrophil behaviors in the brain (42). We further quantified a “hypersegmented” population of neutrophils at the 24-hr time point, with increased transmigratory potential to characterize the dynamics of neutrophilia during this period of early inflammatory linkage between the lung and brain (44).

The persistence of neutrophil engagement at and ingress across the BBB up to +72 hrs after the third i.n. LPS exposure led us to then examine other features of inflammatory activation at the neurovasculature through this observational window. For example, the response to neutrophilia can involve the acute phase protein fibrinogen, typically as a protective response to NETosis that occurs with neutrophil cell death (52). Soluble fibrinogen circulates in plasma, but can deposit in the brain at paravascular spaces, similar to early lesions in multiple sclerosis, where it can attach to axonal nodes and co-localize with microglia (51, 56). Paravascular fibrinogen can also serve as sites for neutrophil adhesion in Alzheimer’s related models (41). Interestingly, while fibrinogen accumulation at the endothelium, as well as paravascularly, followed a similar trend to neutrophil engagement and ingress, synaptic targeting by fibrinogen, with increasing loss of synaptic densities did not persist up to the +72 hrs timepoint. Rather, the amount of fibrinogen-labeled Homer-1 densities peaked between +6 and +24 hrs, with loss of synaptic contact happening primarily within the +6 hr endpoint. This suggests that while persistent fibrinogen deposition may serve to aid continued adhesion and ingress of neutrophils through the +72 hr endpoint, the accumulation of neutrophils may not support continued depletion of synaptic structures.

To corroborate this, we screened for changes in IBA1 and CD68 expression as proxies for typical microglial activation states in early acute inflammatory stages (i.e., microgliosis and enhanced phagocytic activity). The trends of both markers followed that of fibrinogen targeting Homer-1 synaptic densities, peaking between +6 and +24 hrs after the third i.n. LPS exposure. These results appeared incongruous with the rapid depletion of synapses at +6 hrs towards an asymptotic boundary sustained through the +72 hr endpoint; a continued decline in synaptic content is to be expected given the microglial activation and fibrinogen binding timelines. We hypothesized that the persistent neutrophilia in the parenchyma was leading to neutrophil-microglia interactions that biased microglia towards a less phagocytic phenotype between the +6 and +24 hr endpoints that resulted in measurable differences in cellular protein levels between the +24 and +72 hr endpoints (61–64). Using detected objects that were tdTomato Ly6G+ and IBA1+ as a proxy for neutrophil-microglia interactions, we identified a significant fraction of extravasated neutrophils were in contact with microglia by the +6 hr endpoint, that sustained throughout the +72 hr time course. Together, results imply neutrophil-dependent tuning of microglial activation profiles that preferentially downregulates the phagocytic potential of microglia during early induction of lung-brain immune coupling by three subacute exposures to i.n. LPS

Notably, sustained elevated levels of fibrinogen through the +72 hr endpoint, suggests potential impairment of fibrinolytic mechanisms driven by high levels of neutrophil engagement at the BBB (53, 66). We corroborated this in Supplemental Figure S1 by assessing changes in co-expression of VCAM-1 and glucose transporter 1 (GLUT-1) because of known regulatory roles for GLUT-1 in the structural integrity of endothelial function (67–69). We first confirmed that GLUT-1 expression remained constant across the three terminal endpoints compared to saline control (Supplemental Figure S1A). As a result, we interpret that the consistency of VCAM-1 and GLUT-1 co-expression (Supplemental Figure S1B) with the VCAM-1+ CD31+ double immunoreactivity (Figure 3H) and dextran extravasation (Supplemental Figure S2A) reflect an endothelial cell inflammatory profile with potentially GLUT-1-independent mechanisms for changes in barrier permeability that contribute to fibrinogen deposition. This phenomenon may be explained by dynamic flux between competing upregulation and downregulation pathways from neuroinflammation and hypoxia from ALI, respectively (68–70). While deciphering the exact mechanisms governing GLUT-1 expression levels are outside the central focus of this study, the stable expression of GLUT-1 expression is consistent with previous observations of the role the glucose transporter plays in coordinating pro-inflammatory phenotypes in both microglia and neutrophils (67, 71); and is consistent with our observations of individual and joint activation in both of these cell types. Moreover, while previous work has demonstrated activation through the OB, we do not observe vascular vulnerabilities in the vessels of the OB at the endpoints evaluated in our study. While activation of the OB may occur during the initial exposure, it does not seem to be sufficient to drive significant inflammation in other brain regions without concomitant lung injury, as demonstrated by the BAL and qPCR data in Figure 1.

Lastly, reverse-translated our findings of neutrophilia and endothelial dysfunction to in vitro MPS’s that model the BBB. In these MPS translational experiments, we corroborated our findings of increased potentiation of neutrophils for BBB adhesion and transmigration with hallmarks of neutrophil inflammatory activity. The murine MPS, additionally, allowed us to corroborate the sex-independence observed in our flow cytometry analysis. Given the similarities in results of the murine and human MPS models, we posit that our proposed neutrophil-centric paradigm for lung-to-brain neuroimmune linkage is likely sex-independent in humans as well. Further epidemiologic and neuropathologic studies are needed to verify this observation of sex-independence.

Using our three-dose subacute exposure paradigm as an antecedent to pathologic neuroimmune activation from chronic exposure, our results in aggregate implicate cotemporaneous neutrophilia in the lung, circulation, and cerebrovasculature as a key early mediator of lung-brain inflammatory coupling. Particularly, the downregulation in the phagocytic ability of microglia within +72 hrs after three i.n. exposures may provide insight into the correlations between increased risk for neurodegenerative diseases (such as Alzheimer’s and Parkinson’s Diseases) and the frequency of exposure to poor air quality containing aerosolized inflammatory factors (3, 9, 28–30). It is possible that continued inflammatory exposure with ensuing neutrophil recruitment to the parenchyma may lead to terminally immunosuppressed microglia with decreased phagocytic potential, thereby leaving the brain vulnerable to the accumulation of waste factors, such as aggregates of misfolded amyloid-β and α-synuclein, relevant the Alzheimer’s Disease and Parkinson’s Disease, respectively (60, 62, 63). Our work identifies a potential neutrophil-driven mechanism in early stages of activations along the lung-brain neuroimmune axis relevant to better understanding how chronic respiratory exposure increases risk for pathologic brain disorders. Continued insights from such studies may help identify therapeutic targets to mitigate neuroinflammatory responses and is the subject of our ongoing research.

## Methods

### Sex as a biological variable

In the in vivo mouse experiments and in vitro murine MPS samples, both male and female sexes were accounted for as a variable. An initial screen of BAL and brain transcripts involved on male mice, but all subsequent experiments based on questions from the results of the initial screen validated that similar effects were observed in both male and female mice before aggregating them. The in vitro murine MPS uses neutrophils isolated from both male and female mice. The human MPS system only included the male sex as a biological variable due to the origin of the iPSC donor source and the human volunteer for neutrophils in circulation.

### Mice

All mice were housed under a 12-hour light/dark cycle with access to water and regular chow ad libitum in temperature and humidity-controlled environment. Wild-type male and female C57BL/6J were purchased from The Jackson Laboratory (Bar Harbor, ME, USA) and housed at Duke University and the University of Rochester Medical Center (URMC). Male and female Catchup mice (C57BL/6-*Ly6g*(tm2621(Cre-tdTomato)Arte)) were bred and maintained by the Gelbard laboratory at URMC. The Catchup mice were genotyped following previously reported procedures (39). In brief, ear biopsies were collected around weaning age (3-4 weeks) and digested using a Mouse Direct PCR Kit (Fisher; cat: B40015) following the manufacturer’s protocols. The forward primers for the wild-type (WT) and knock-in (KI) are 5’-GGT-TTT-ATC-TGT-GCA-GCC-C-3’ and 5’-ACG-TCC-AGA-CAC-AGC-ATA-GG-3’, respectively, with a common reverse primer of 5’-GAG-GTC-CAA-GAG-ACT-TTC-TGG-3’. A control primer for separate gene was used consisting of a forward 5’-GAG-ACT-CTG-GCT-ACT-CAT-CC-3’ and reverse 5’-CCT-TCA-GCA-AGA-GCT-GGG-GAC-3’.

### Reagents and Antibodies

All reagents used in this manuscript are noted in line with manufacturer details where appropriate in the methods. We have also compiled this information in Tables S1-S3.

### Delivery of LPS

Mice that received i.n. delivery of LPS (*E. coli* O111:B4; Sigma; cat: L2630) were dosed at a concentration of 0.25 μg/μL diluted in 0.9% NaCl saline solution per delivery timepoint at either single timepoint (10 μg total LPS), or across three timepoints spaced 24 hours apart across a 48-hour treatment period (0 hrs, 24hrs, 48hrs; 30 μg total LPS). Mice that received i.p. delivery of LPS which was performed under isoflurane (Patterson Veterinary, Greeley, CO) anesthesia. All mice recovered from intranasal LPS instillation and were included in the study.

### Collection and analysis of bronchoalveolar lavage fluid (BALF)

Bronchoalveolar lavage (BAL) samples were collected 24 hours after the final i.n. exposure in the assigned treatment group. BAL was performed by cannulation of the trachea and gentle instillation/aspiration (3 times) of 1.0 ml of PBS with protease inhibitor cocktail tablets (Roche, Indianapolis, IN). The lavage fluid was centrifuged at 4000 rpm for 5 minutes, and the supernatant was stored at -80 °C for later protein content assessment. The cell pellet was treated with red-blood-cell lysing buffer (BD Biosciences, San Diego, CA), washed, and resuspended in 200 µl of PBS. Total cell counts were determined with a hematology analyzer (Scil Vet ABC, Gurnee, IL), centrifugated onto cytoslides (Cytospin 3, Shandon Inc, Pittsburg, PA) and stained with Diff-Quick (Dade-Behring Inc., Newark, DE). Differential cell counts were obtained by microscopic counting of a minimum of 200 cells/slide, using standard morphological and staining criteria. The total protein in the BALF was measured with a Qubit 3.0 Fluorometer (Invitrogen, Waltham, MA).

### Real-Time PCR (qPCR)

Brains were harvested from wild-type C57BL/6J mice 24 hours after the final i.n. dose of their assigned treatment group. Brain tissue was homogenized in a NextAdvance bullet blender at 4 °C in Eppendorf Safe-Lock tubes, followed by extraction of total RNA from the homogenates using the RNeasy Lipid Tissue Mini Kit from Qiagen (Germantown, MD, USA) following the manufacturer’s instructions. cDNA synthesis was performed right after the RNA isolation with the High-Capacity RNA-to-cDNA Kit (Applied Biosystems, Foster City, CA) following the manufacturer’s instructions, using 200 ng of RNA per 20 μl cDNA reaction. The resultant cDNA was analyzed using TaqMan® Gene Expression Assays (Applied Biosystems, Foster City, CA, USA) in 10 μl reactions. Thermal cycling was performed with the Real-Time PCR system QuantStudio 5 (AppliedBiosystems). Each sample was performed in triplicates and normalized to the endogenous *Act-b* gene expression. The CT value of each well was determined using the QuantStudio 5 software and relative quantification was determined by the ΔΔCT method. Specific TaqMan® assays used are listed in Table S4.

### Flow Cytometry

Cell pellets were resuspended in FACS buffer (1% BSA + 2.5 mM EDTA in 1x PBS), and Trypan Blue (ThermoFisher; cat: 15250061) stained aliquots were counted on a hemacytometer. After counting, aliquots of 5E6 cells were first blocked with 20 μg of Fc block (ThermoFisher; cat: 14-9161-73) for 30 minutes on ice. The cells were pelleted at 300 xg for 5 minutes at 4 °C and resuspended in FACS buffer with fluorophore-conjugated primary antibodies (CD11b PE-TexasRed, ThermoFisher, cat: RM2817; Ly-6G/Ly-6C, AlexaFluor700, cat: 56-5931-82) validated for flow cytometry. Staining reactions were performed in the dark at 4 °C with buffers and reagents kept on ice until use. The cells were pelleted at 300 xg for 5 minutes at 4 °C and washed via resuspension in FACS buffer and centrifugation at the same conditions. The cells were then fixed via resuspension in 100 μL of 2% PFA in FACS buffer and incubated in the dark at room temperature for 10 minutes. The cells were then pelleted and resuspended in 500 μL of FACS buffer. Samples were run on a FACSymphony A1^TM^ (BD Biosciences) controlled by FACS Diva software (BD Biosciences). Results were processed using FCS Express^TM^ (De Novo Software). Details for peripheral blood collection and sample handling can be found in the Supplementary Materials.

### Two-photon intravital microscopy

2P intravital microscopy was performed on Catchup mice with somatosensory cortical cranial windows implanted at least 30 days prior to induction of assigned i.n. exposure group. Full details on cranial window surgery can be found in the Supplementary Materials.

An Olympus Fluoview FVMPE-RS Twin-Laser Gantry multiphoton microscope system equipped with two tunable lasers (Spectra-Physics; MaiTai HP DeepSee and InsightX3) was used for intravital imaging. Two-photon excitation was achieved using 100 fsec laser pulses (80 MHz) tuned to 840 nm with a power of ∼50 mW measured at the sample focal plane. For anesthetized imaging sessions, mice were anesthetized via i.p. delivery of a fentanyl cocktail prepared with the same protocol used for cranial window implantation. To visualize the brain vasculature, 100 μL of high molecular weight (2,000 kDa) FITC-Dextran was introduced via tail vein injection at a concentration of 50 mg/mL in 0.9% NaCl saline solution 5 min before intravital imaging. Rodent body temperature was maintained at 37°C with a heating pad, and the animal’s eyes were protected with lubricant ointment. Intravital imaging was performed using 4x digital zoom at 512 x 512-pixel resolution. Stacks containing 41 slices (1 mm step size) were imaged every 90 seconds for 15 min to obtain 4D (x, y, z, t) “hyperstack” data used for analysis using ImageJ. Full details available in the Supplementary Materials.

### Immunofluorescence

IHC staining of paraformaldehyde (PFA)-fixed brain slices (40 μm thick) was performed as previously reported (72). Sections were washed to remove cryoprotectant and reduce autofluorescence. Sections were subsequently incubated in blocking buffer (1.5% BSA, 3% normal goat serum, 0.5% Triton-X, and 1.8% NaCl in 1X PBS) containing primary antibodies at room temperature overnight. After washing, sections were then incubated overnight in Alexa Fluor conjugated secondary antibodies (ThermoFisher 1:750) in the same blocking buffer composition. Finally, sections were washed three times with 1X PBS + 1.8% NaCl, mounted on glass slides with Prolong Diamond Antifade Reagent (Invitrogen P36961). For dextran leakage from small and large vessels 40 kDa FITC-Dextran (ThermoFisher; cat: D1845) was injected 5 minutes prior to perfusion. Full details of the IHC staining protocol can be found in the Supplementary Materials.

Immunocytochemistry of the murine MPS were stained similarly to previous protocols (73), where the cells were briefly rinsed with ice-cold 1x DPBS and then fixed for 12 minutes at room temperature with 4% PFA in 1x PBS. The cells were washed, blocked with 5% BSA, and stained via an indirect (primary and secondary) immunolabeling protocol before the MPS was disassembled and the membranes mounted onto coverglass for imaging. Full details of the murine MPS staining protocol can be found in the Supplementary Materials.

Both IHC and ICC prepared samples were imaged on the “grid-confocal” structured illumination acquisition system detailed in the Supplementary Materials.

### Image analysis

ICC and IHC images were analyzed for semi-quantitative analysis using Volocity 3DM Quantitation Software (Quorum Technologies). A summarized workflow includes an initial illumination correction and “fine noise filter” over all images. Immunoreactive objects were determined as signal at least n standard deviations above the mean intensity in each channel. Intensity, volume, and position details were summarized in the software and exported to Excel for further workup before importing the data into GraphPad Prism for statistical analysis. Image panels were assembled using Affinity Designer. Full details are available in the Supplementary Materials.

### Assembly of Microphysiological Systems

Model murine and human MPSs were assembled following previously published protocols of culturing human-derived cells on μSiM devices in either an open well or flow cell configuration (48, 74–76). The murine MPS experiments were conducted using an open well configuration. The human MPS experiments were conducted using the flow cell configuration. Full details of μSiM assembly can be found in the Supplementary Materials.

### Neutrophil Engagement and Activation in Murine MPS

The murine MPS models were constructed by culturing immortalized murine brain endothelial cell line (bEnd.3) purchased from ATCC (cat: CRL-2299) on μSiM devices. All MPS maintenance and exposure periods to treatments were conducted in an incubator with 5% CO_2_ at 37 °C. Full details of bEnd.3 cell line maintenance and murine MPS setup can be found in the Supplementary Materials.

The bEnd.3 monolayers in the murine MPS were stimulated with CytoMix, a cytokine cocktail comprised of 150 ng/mL IL-1β (PeproTech; cat: 211-11B) and 150 ng/mL TNFα (PeproTech; cat: 315-01A) in Dulbecco’s Modified Eagle Medium (ThermoFisher; cat: 10567022) + 1% fetal bovine serum (Atlas Biologicals; cat: F-0500-D). Prior to stimulation, the cells were first serum deprived in this 1% serum composition without CytoMix overnight. After 6 hours of stimulation, freshly isolated neutrophils from “vehicle” and “ALI” C57BL/6J mice of either sex were added to the MPS and incubated for 3 hours before fixation, immunocytochemical staining, and imaging. Approximately 175,000 neutrophils in 75 μL were added to each open well murine MPS. Full experimental details can be found in the Supplementary Materials.

### Neutrophil Transmigration in Human MPS

Human MPS models were constructed using extended endothelial cell culture method-brain microvascular endothelial cell like cells (EECM-BMECs) differentiated from IMR90-4 hiPSCs (WiCell; cat: iPS(IMR90)-4) and cultured in μSIM devices as previously described (48, 74–76). Full details of the culturing the human MPS models can be found in the Supplementary Materials.

Neutrophil trafficking studies were performed using previously described methods (77, 78). Briefly, whole blood was drawn from consenting donors and neutrophils were subsequently isolated using a density separation gradient (1-Step Polymorphs, Accurate Chemical & Scientific Co., Westbury, NY). Following isolation, neutrophils were used within three hours to ensure viability. For experimentation, freshly isolated neutrophils were incorporated into hESCR media at a concentration of 3 million cells/mL, deposited into µSiM devices with EECM-BMECs, and placed into an incubation stage (Okolab s.r.l., Pozzuoli, Italy) set to a physiological temperature of 37 °C. The stage was part of a phase-contrast microscopy system (Nikon Ti2E Inverted Microscope, Nikon Corporation, Tokyo, Japan) and transmigration studies were recorded for 30 minutes at a framerate of 0.25 Hz with a 40x long working distance lens (NA 0.55). Following recording, neutrophil transmigration at *t* = 0 and *t* = 30 minutes was characterized for control and treated devices respectively via manual counting. Additionally, neutrophil association with the endothelial barrier was assessed at *t* = 30 minutes via manual counting. Statistical comparisons were made with a two-way ANOVA with Tukey’s multiple comparison correction and unpaired t-test respectively.

### Statistical Analysis

One- and two-way analysis of variance (ANOVA) with Holm-Sidak’s or Tukey’s post-hoc corrections for multiple comparison tests were used to analyze all data. The specific chosen analysis and post-hoc corrections for the independent data sets are identified, with accompanying rationale, in the appropriate figure legends. All data are presented as mean ± standard error (SEM) with significance at p ≤ 0.05, and all data points representing biological replicates plotted. The sample sizes are identified in the figure legends for each respective dataset.

## List of Abbreviations

2PM: two-photon microscopy
BAL: bronchoalveolar lavage
BBB: blood-brain barrier
CD68: Cluster of Differentiation 68
CNS: central nervous system
EECM-BMEC: extended endothelial cell culture method-brain microvascular endothelial-like cells
GLUT-1: glucose transporter 1
H3-cit: citruillinated histone H3
hiPSCs: human induced pluripotent stem cells
i.n.: intranasal
IL1β: interleukin 1β
LPS: lipopolysaccharide
Ly6G: lymphocyte antigen 6 complex 6GD
MCP1: monocyte chemoattractant protein 1
MPO: myeloperoxidase
MPS: microphysiological system
µSiMs: optically transparent nanoporous silicon membranes
NLR: neutrophil to lymphocyte ratio
NVU: neurovascular unit
OB: olfactory bulb
PECAM1: platelet endothelial cell adhesion molecule 1
PSD95: post-synaptic density 95
SLM: stratum lacunosum moleculare
TLR4: toll-like receptor 4
TNFα: tumor necrosis factor α
VCAM1: vascular cell adhesion molecule 1

## Supplementary Materials

Extended methods with information on all key reagents tabulated can be found in the supplementary document. Additional processed figures can also be found in the supplementary document. Sample 2P videos of neutrophil behavior in murine cerebrovasculature under ALI and vehicle conditions are as follows: Supplemental Video 1 – stalling, crawling, and transmigration in ALI condition; Supplemental Video 2 – stalling in ALI condition; Supplemental Video 3 – minimal evidence of neutrophils in vehicle condition (male); Supplemental Video 4 – female vehicle condition.

## Declarations

### Ethics of Approval and Consent to Participate

All procedures involving mice were performed under strict compliance to protocols approved by the Institutional Animal Care and Use Committees at both Duke University Medical Center (DUMC) and University of Rochester Medical Center (URMC) and developed in accordance with the National Research Council Guide for the Care and Use of Laboratory Animals, 8^th^ edition (79). Duke University and the University of Rochester Medical Center are AAALAC-accredited institutions.

The use of human iPSCs (WiCell; cat: iPS(IMR90)-4) and human neutrophils isolated from whole blood collected from consenting donors was conducted under strict adherence to a protocol approved by the Institutional Review Board at the University of Rochester (protocol #00004777).

### Data and Materials Availability

All tabulated data involved in presented figures are included in a supplementary data file. Data files, sample images, figures, and supplementary videos will also be made accessible in an online repository (https://osf.io/28f9a/?view_only=d1c6f7c21ae74ed5906626c9d708ea14) that will be kept up to date.

## Author Contributions

NT, HAG, HL, and WC conceptualized the study. NT, HAG, AKM, and JLM provided funding, supervision, and resources. HL, WC, LL, JD, AIC, KSS, AP, CL, and DA all conducted experiments that resulted in contributory inclusion into the presented figures. HL, WC, LL, JD, AIC, KSS, DA, JLM, AKM, NT, HAG contributed to the design and optimization of methodology used in this work. WC, HL, LL, DA analyzed the data. WC and HL generated the figures. WC, HL, MM, and KSS contributed to writing of the manuscript. WC, HL, MM, KSS, NT, and HAG were involved in the review and editing of the manuscript. WC and HL are listed as co-first authors due to HL initializing the study and WC performing key final experiments and overall analyses of data that resulted in the framing of the paper. The order of these two authors was based on last name.

## Acknowledgments

We would like to acknowledge the assistance and resources used by the URMC Center for Advanced Light Microscopy and Nanoscopy (CALMN). Vector graphics used in the diagrams were sourced from online repositories with open-source agreements. The accreditations for the various graphics are as follows: mouse icon (Figures 1, 6) by Ben-Murrell on Bioicons.com is licensed under CC0; immune cell icons (Figures 1, 2, 4, 6, 7) by Servier on Bioicons.com is licensed under CC-BY 3.0 Unported; human diagram (Figures 1, 2, 7) icons source from Flaticon.com; alveolar health, blood brain barrier, and microglia at synapse icons (Figures 1, 2,4) are open source vectorial images taken from sciencefigures.org under Open Design License 1.1.

