## Supplemental Document for "Dynamics of Neutrophilia at the Neurovascular Unit Arising from Repeated Pulmonary Inflammation"

**A Neutrophil-Centric Lung-to-Brain Neuroimmune Axis in a Murine Model of Repeated Respiratory Inflammation**

### Extended methods

#### Peripheral Blood Isolation via Cardiac Puncture

Blood samples for flow cytometry and in vitro neutrophil studies in the murine microphysiological system (MPS) were collected via cardiac puncture after mice were deeply anesthetized using a ketamine/xylazine cocktail (90 mg/kg ketamine, 10 mg/kg xylazine in 0.9% NaCl saline solution). The collected blood was transferred into 1.5 mL microcentrifuge tubes with 50 uL of EDTA starting volume. The collected blood was first incubated in 2x volume of ACK Lysing Buffer (ThermoFisher; cat: A1049201) for 3 minutes at room temperature. The leukocyte-enriched cell suspension was pelleted via centrifugation at 300 xg for 5 minutes at room temperature and the resulting supernatant removed via aspiration. The pellet was resuspended in ice-cold buffer (1x PBS + 0.5% BSA + 2 mM EDTA) and pelleted via centrifugation at 300 xg for 5 minutes at 4 ^o^C. The pellet was then used for either flow cytometry or neutrophil isolation.

#### Murine Neutrophil Isolation

Both vehicle and lipopolysaccharide (LPS; strain *E. Coli* O111:B4)-stimulated C57BL/6 mice of both sexes were used for isolation of neutrophils from the peripheral blood to reflect healthy “veh Neuts” and ALI-activated “ALI Neuts” populations. Lymphocyte-enriched cell pellets were obtained following the “Peripheral Blood Isolation via Cardiac Puncture” protocol above. This pellet was resuspended in 20 uL of antibody cocktail from the MojoSort^TM^ Mouse Neutrophil Isolation Kit (BioLegend; cat: 480058), diluted with 200 uL of the same buffer used in the previous step, and left on ice for 15 min. The cell suspension was then diluted to 4 mL total volume in the buffer and centrifuged at 300 xg for 5 minutes at 4 ^o^C. The supernatant was aspirated, and the pellet resuspended in 20 uL of nanobeads from the isolation kit, diluted with 200 uL of buffer, and incubated on ice for 15 min. The labeled cell suspension was pelleted again under the same conditions and the pellet resuspended in 2 mL of fresh buffer and placed on a magnetic column for 5 minutes. The free supernatant was transferred to a fresh tube, and the magnetic bead pellet was used to repeat this step once more with the free supernatant collected into the same tube. The final 4 mL supernatant was placed on the magnet for 5 minutes to capture any residual magnetic beads and transfer the neutrophil-enriched supernatant to a final collection tube. The cell suspension was pelleted via centrifugation at the same conditions and resuspended in 500 uL of cell culture media. A cell count was performed by diluting an aliquot of the neutrophil suspension in Trypan Blue (ThermoFisher; cat: 15250061). The cells were then diluted to the final concentrations used in the study.

#### Cranial Window Surgery

Catchup (C57BL/6-*Ly6g*(tm2621(Cre-tdTomato)Arte)) mice were anesthetized via i.p. delivery of a fentanyl cocktail (0.05 mg/kg fentanyl, 5.0 mg/kg midazolam, 0.5 mg/kg dexmedetomidine in 0.9% NaCl saline solution) during the cranial window implantation surgical procedure. Body temperature was maintained at 37°C with a heating pad, and the animal’s eyes were protected with ophthalmic lubricant ointment. All surgical procedures adhered to the aseptic technique. Cranial window implantation surgeries were performed in the Majewska laboratory as described (*1, 2*). Mice were fixed in a stereotaxic frame, hair was removed with Nair hair removal cream, the dermis was sterilized with three alternating applications of 70% ethanol and beta-iodine (i.e., povidone-iodine), and the skull was exposed through a scalp incision. A 3-mm biopsy punch (Integra) was then used to create a circular score on the skull over the primary somatosensory cortex (S1). A 0.5 mm drill bit (Fine Science Tools, Foster City, CA) was used to drill through the skull for the craniotomy, tracing the 3-mm score. A 5-mm coverslip attached to a 3-mm coverslip (Warner Instruments) by UV glue (Norland Optical Adhesive, Norland) was then slowly lowered into the craniotomy (3-mm side down). The coverslip was carefully secured with C&B Metabond dental cement (Parkell). A custom head plate produced by eMachineShop® (Mahwah, NJ, USA) using designs courtesy of the Mriganka Sur laboratory (Massachusetts Institute of Technology) was then secured onto the skull with the same dental cement. Any exposed skull and the incision site were further covered and sealed with dental cement. Mice were administered the analgesics slow-release buprenorphine (5 mg per kg subcutaneously for 72 h duration) and carprofen (5 mg/kg, i.p. every 24 h) and monitored for 72 h postoperatively. Mice were allowed to recover for a minimum of 30 days and a maximum of 2 months before imaging to reduce parenchymal inflammation and minimize bone regrowth.

#### Processing and Analysis of 2P Intravital Microscopy Images

“Hyperstack” data containing changes in intensity data encoded over spatial coordinates and time (x, y, z, t) were imported into ImageJ for each channel. The “true” signal in each channel were identified via standard intensity thresholding to produce binary image stacks. The threshold method for each stack was selected after determination of the optimal method from the “Try all” parameter in the “Auto Threshold” tool. Binarized images were used as masks to represent spatial positions of where the GLUT1+ vessels were as well as flux of FITC-Dextran. Co-occurrence of the two was determined by multiplying the two masks to find double positive voxels at each time point. The sum of individual and “colocalized” voxel counts were tabulated and exported as .csv files for downstream processing in Excel with statistical analysis in GraphPad Prism.

#### µSiM Assembly

Ultrathin (<100 nm thick) silicon nitride membranes (NPSN100-1L, SiMPore Inc, West Henrietta, NY) with nanoscale sized pores (~60 nm size, ~15% porosity) were incorporated into µSiM devices and used for endothelial cell culture as a scaffold. Assembly instructions for the µSiM device have been described in detail previously to create open well (*3, 4*) and flow cell configurations (*5, 6*). The open well devices were used for staining and flow cells were used for neutrophil studies. For both devices, constituent components were irreversibly bonded around silicon nitride membranes using pressure sensitive adhesive (PSA). To ensure sterility, devices were made in a biosafety cabinet and sterilized with UV light for 20 minutes prior to use.

#### Murine Microphysiological System

The murine MPS models were constructed by culturing immortalized murine brain endothelial cell line (bEnd.3) purchased from ATCC (cat: CRL-2299) and maintained according to manufacturer protocols. Specifically, the bEnd.3 cells between passage numbers 24-30 were maintained in a flask containing Dulbeco’s Modified Eagle Medium (DMEM, ThermoFisher; cat: 10567022) supplemented with 10% Fetal Bovine Serum (FBS, Atlas Biologicals; cat: F-0500-D) and passaged with 0.25% trypsin-EDTA (ThermoFisher; cat: 25200056) when the cells are > 90% confluent. Prior to addition of bEnd.3 cells, the assembled μSiM devices were plasma-cleaned for 10 minutes, followed by brief UV irradiation, and finally coated with 0.1 mg/mL poly-D-lysine (PDL; Sigma; cat: P1149) on the exposed layer of the silicon nitride membrane. The coated μSiM devices were then seeded with bEnd.3 cells at a density of 2k cells/device and given fresh media after 3 hrs, followed by media replenishment every 24 hours until the barriers looked visually confluent (> 90%) under an optical microscope and were ready for experiments.

#### Human Microphysiological System

Human MPS models were constructed using extended endothelial cell culture method-brain microvascular endothelial cell like cells (EECM-BMECs) differentiated from IMR90-4 hiPSCs and cultured in μSIM devices as previously described (*3, 4, 7, 8*). Briefly, μSiMs were coated with 400 μg/mL collagen IV (Sigma; cat: C5533) and 100 μg/mL fibronectin (Sigma; cat: F1141) prior to seeding EECM-BMECs at a density of 40,000-50,000 cells/cm^2^ in hECSR media. hECSR media is comprised of human endothelial serum-free media (ThermoFisher; cat: 11111-044) supplemented with 2% v/v serum-free B-27 supplement (ThermoFisher; cat: 17504044) and 20 ng/mL human fibroblast growth factor 2 (Tocris; cat: 233-FB-500). After 2 h, the media was replaced with fresh hECSR media and replenished every 24 h until day 7 when the human MPS devices were used for neutrophil migration studies.

#### Immunohistochemistry

Mice were deeply anesthetized with a cocktail of 100 mg/kg ketamine and 10 mg/kg xylazine in saline solution and transcardially perfused with PBS and 4% paraformaldehyde (PFA). Brains were carefully extracted and post-fixed for 24 hr in 4% PFA. Free-floating coronal brain sections (40 μm thickness) were cut using a vibratome and stored in cryoprotectant (30% PEG300, 30% glycerol, 20% 0.1 M phosphate buffer, and 20% ddH2O) at -20˚ C. Cryopreserved sections were washed three times in 1X PBS followed by another wash in 0.1 M glycine in 1X PBS (to reduce autofluorescence). Sections were subsequently incubated in blocking buffer (1.5% BSA, 3% normal goat serum, 0.5% Triton-X, and 1.8% NaCl in 1X PBS) containing primary antibodies (VCAM1 MilliporeSigma MAB1398Z 1:200; Iba1 Wako PTR2404 1:1000; CD68 Serotec MCA1957GA 1:1000; Homer1 SynapticSystems 160006 1:500; PSD95 NeuroMab 75-028 1:500; CD31 MilliporeSigma MAB1398Z 1:250; Fibrinogen Dako A0080 1:200 in blocking buffer (1.5% BSA, 3% normal goat serum, 0.5% Triton-X, and 1.8% NaCl in 1X PBS) overnight at room temperature. Sections were washed three times in 1X PBS containing 1.8% NaCl before incubating in solutions of Alexa Fluor conjugated secondary antibodies (ThermoFisher 1:750) in the same blocking buffer composition overnight. Finally, sections were washed three times with 1X PBS+1.8% NaCl, mounted on glass slides with Prolong Diamond Antifade Reagent (Invitrogen P36961). When used, Alexa-633 Hydrazide (ThermoFisher; cat: A30634) was diluted 1:1000 in 1X PBS+1.8% NaCl and applied to sections for 10 min following the first two post-secondary washes; an additional two washes (1X PBS+1.8% NaCl) were performed to rinse the sections of excess Alexa-633 Hydrazide.

#### Immunocytochemistry

The murine MPS were stained using an indirect immunofluorescent labeling protocol. Specifically, cultures were briefly rinsed with ice-cold DPBS, followed by fixation in 4% PFA diluted in a 1x PBS solution for 12 minutes at room temperature with gentle rocking. The fixative was then removed and quenched with 100 mM glycine (BioRad; cat: 1610718) in 1x PBS solution for 5 minutes at room temperature with gentle rocking, followed by two washes with 1x PBS at the same conditions. The cells were then permeabilized using 0.25 % Triton X-100 (Millipore Sigma cat: T9284) in 1x PBS for 15 minutes at room temperature with gentle rocking, followed by one wash with 1x PBS in the same conditions. The cells were then blocked using 5% BSA (Millipore Sigma cat: A3294) in ddH_2_O for 1 hr at room temperature with gentle rocking. This was followed by the primary antibody incubation step rocking overnight at 4 ^o^C, with the primary antibodies diluted in a staining buffer comprised of 2% BSA with 0.01% Triton X-100 in 1x PBS. The following primary antibodies were used – CD106/VCAM-1 (MilliporeSigma; cat: CBL1300; dil 1:250), neutrophil 7/4 or Ly6B.2 (CedarLane; cat: CL8993B; dil 1:200), CD31/PECAM-1 (MilliporeSigma; cat: MAB1398Z; dil 1:250), and citrullinated histone H3 (Novus Biologicals; cat: NB100-57135; dil 1:250). The next day the primary antibody solution was aspirated off, followed by three washes with 1x PBS at room temperature with gentle rocking for 3 to 5 minutes. This was followed by incubation in the secondary antibodies with AlexaFluor conjugates (ThermoFisher) and rocked at room temperature for 1 hr while protected from light (i.e. wrapping with foil). The secondary antibodies were diluted (1:750) in the same staining solution as the primary antibodies were. This was followed by three rounds of washing in 0.1% Tween 20 (MilliporeSigma; cat: P1379) in 1x PBS for 3 to 5 minutes at room temperature, rocking while protected from light. After staining, the coverslips were mounted onto cover glass using ProLong^TM^ Diamond Antifade Mountant with DAPI (ThermoFisher; cat: P36962) and stored protected from light until needed for imaging.

#### “Grid-confocal” Structured Illumination Image Acquisition and Analysis

*Fluorescent Imaging w/ “Grid” Confocal Microscope:* Immunofluorescent microscopy data was acquired using an Olympus BX-51 microscope in “grid-confocal” configuration using an OptiGrid structured illumination optical element (Qioptiq) in the excitation path as previously reported by the Gelbard laboratory (*9*). A Prior Lumen200 illumination source equipped with a Hg lamp (Prior; cat: LM200B1-A) is coupled into the microscope via a liquid light coupler (Prior; cat: LM587). Fluorescent emission from the immunostained samples was detected using a Hamamatsu ORCA-ER scientific camera. The following pairs of excitation and emission filters were used to selectively illuminate and detect biomolecular targets immunolabeled with DAPI, AlexaFluor488, AlexaFluor568, and AlexaFluor647, respectively: 350 nm / DAPI (Semrock; cat: FF02-447/60-25); 405 nm / FITC (Semrock; cat: FF01-524/24-25); 488 nm / TRITC (Semrock; cat: FF01-593/40-25); 568 nm / Cy5 (Semrock; cat: FF01-692/40-25). Volocity 3DM software (Quorum Technologies) was used for hardware integration and control to acquire all images. Synaptic punctae were imaged at 60x magnification (0.3 μm z-step sizes through a 10 μm axial range). Other markers were imaged at 20x and 40x magnification at (1 μm z-steps through a 15 μm axial range). The following infinity-corrected Olympus objectives were used: 20x UPlanApo 0.70 NA, 40x UPlanApo 0.85 NA, or 60x UPlanSApo 1.35 NA oil.

*Volocity Image Analysis:* Immunofluorescent images were analyzed for semi-quantitative analysis using Volocity 3DM Quantitation Software (Quorum Technologies). A fine noise filter was used on all images before applying a set of measurement protocols. The “Find Objects” function was used to identify fluorescent objects above a certain intensity threshold defined by n standard deviations above the average voxel intensity of a given channel, where n ranged from 1.5 to 3 standard deviations depending on the skew of the intensity distributions in each channel. For analysis of synaptic puncta, a reduced ROI of 300x300 voxels from each stack was used to minimize variability due to differences in vascular coverage (i.e., areas without synaptic puncta). Synaptic puncta were identified via a double-positive screen for fluorescent objects that were above the “Find Objects” threshold as identified as local maxima via the “Find Spots” function. These “colocalized” positive voxel objects were then labeled as synaptic puncta for subsequent analysis. Arterial VCAM-1 vessels were defined as VCAM1+ objects touching Alexa Fluor Hydrazide+ objects. Microglial CD68 was quantified CD68+ objects colocalized with IBA1+ objects. Extravascular fibrinogen was quantified as fibrinogen+ objects not in contact with CD31+ objects.

### SUPPLEMENTAL FIGURES

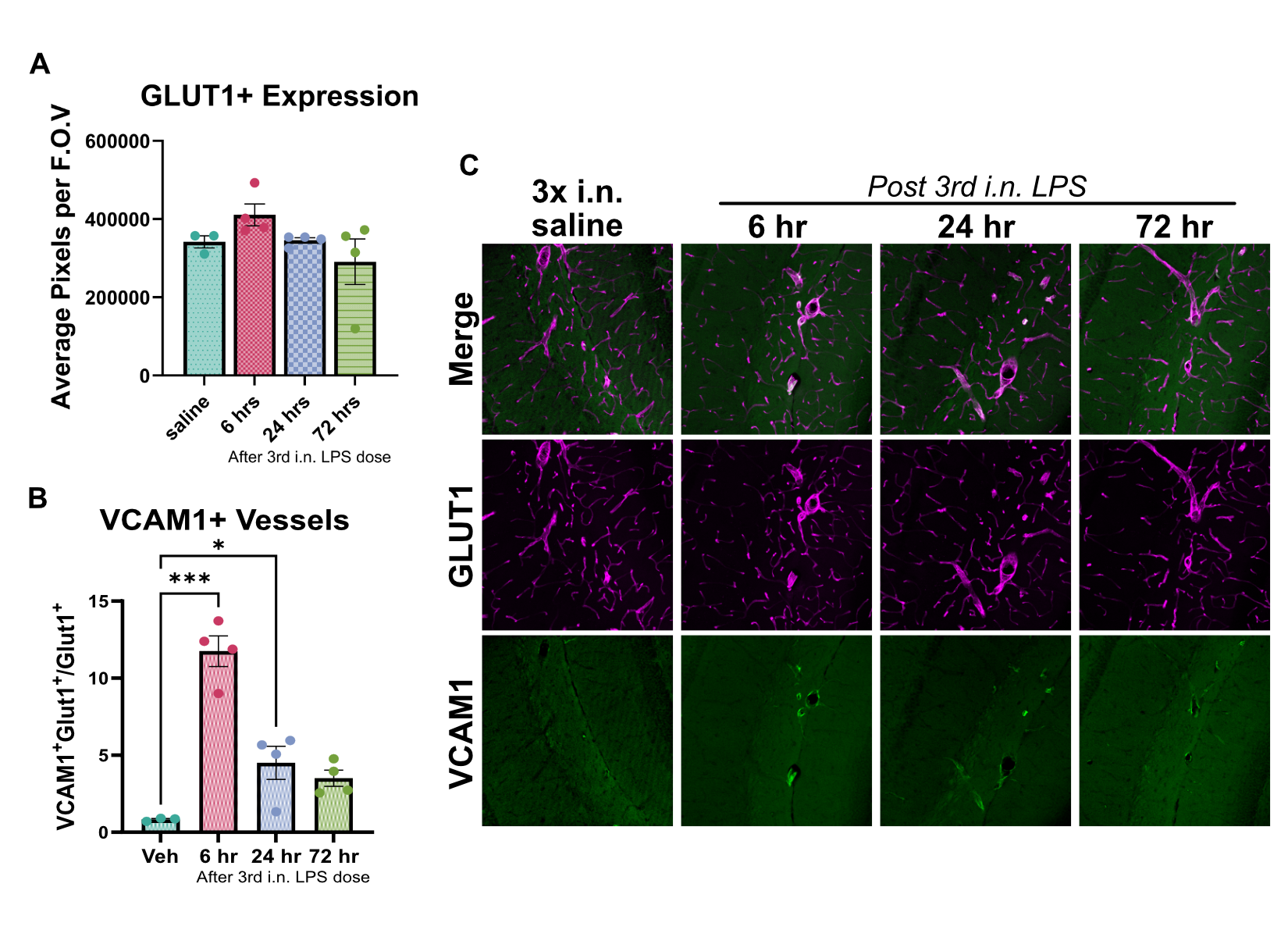

#### Figure S1. BBB Inflammatory activation probed by VCAM1 upregulation in GLUT1+ vessels.

**(A)** GLUT1+ vessels did not change notably in their distribution when thresholded to identify “true” signal from GLUT1 immunoreactive objects. **(B)** VCAM1 immunoreactivity was assessed in GLUT1+ vessels to identify trends in endothelial cell inflammatory activation in the multi-hit i.n. LPS ALI paradigm. **(C)** Representative IHC panels for VCAM1 and GLUT1 staining in harvested brain tissue sections.

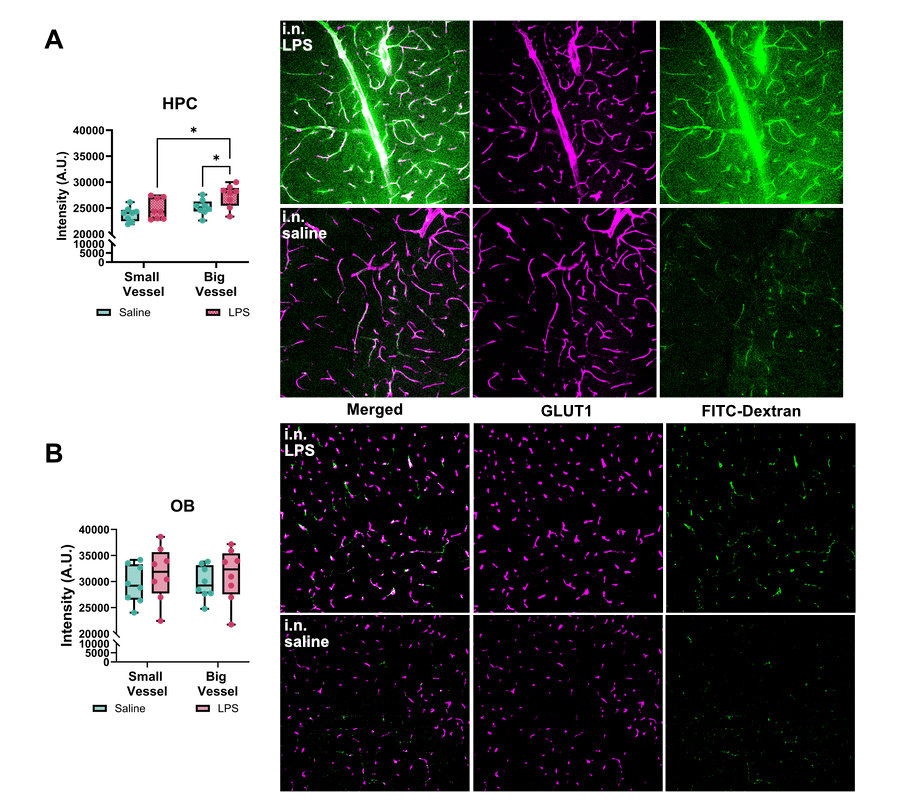

#### Figure S2. FITC-Dextran leakage enhanced in big vessels of HPC after multi-hit i.n. LPS ALI paradigm.

**(A)** Colocalization analysis of FITC-Dextran 40 kDa and GLUT1 in the vessels of the HPC reveal increased leakage from big vessels, but not smaller vessels. This is represented by vessels with much thicker GLUT1 staining that exhibit a “smearing” or “bleeding” of FITC-Dextran fluorescence beyond the boundaries of the vessel. **(B)** Similar examination of the OB does not demonstrate appreciable differences in FITC-Dextran 40 kDa leakage from either vessel types.

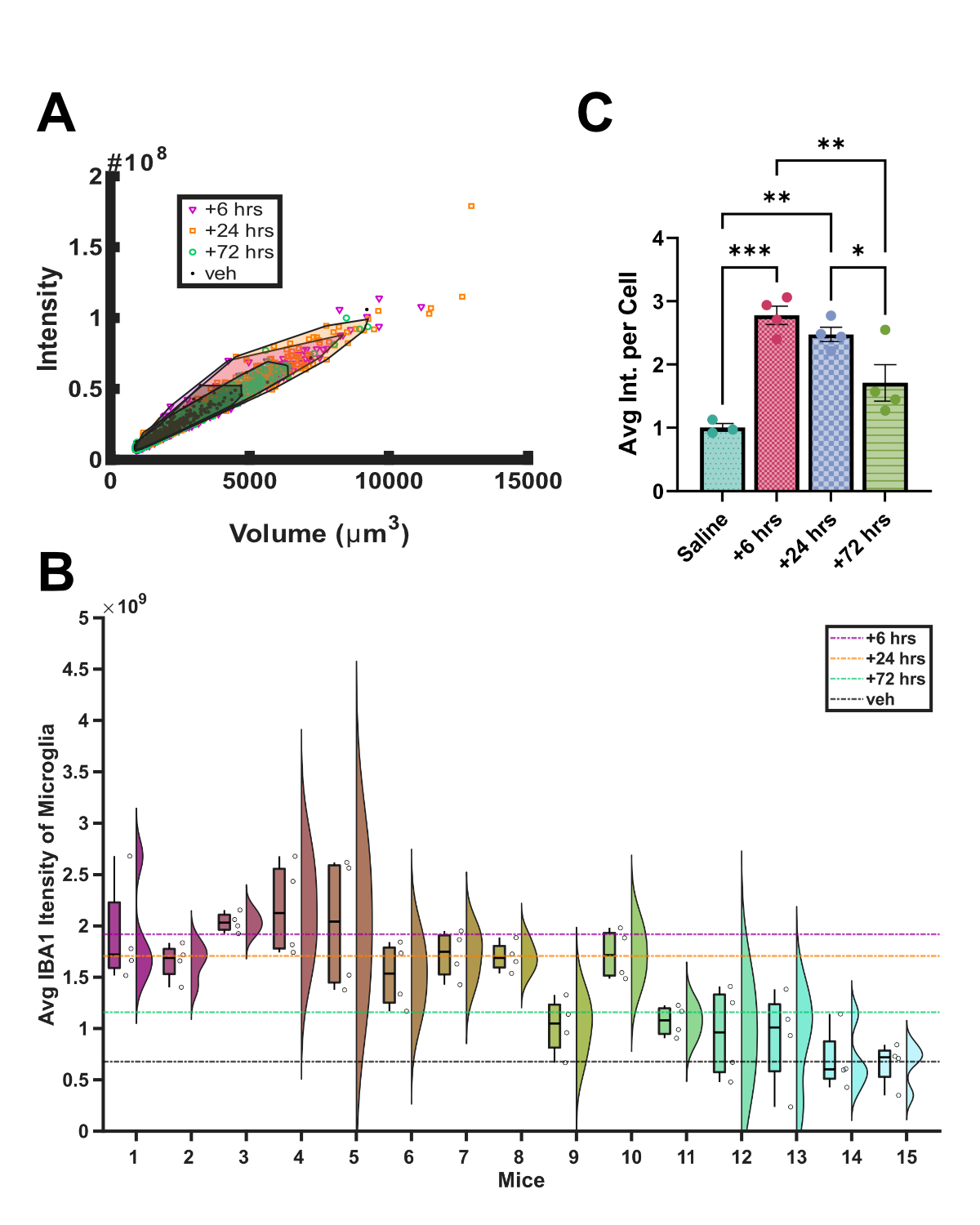

#### Figure S3. Corroborating ensemble aggregated changes in IBA1+ microglial objects in segmented single cell distributions.

**(A)** Plotting of individual IBA1+ microglia segmented from IHC micrographs used in Figure 7. The distribution of points and the enclosed area demonstrate increased microglial activations at an individual cell level at +6 hrs (magenta) and +24 hrs (orange) that is downregulated by +72 hrs (green) when compared to the vehicle group (gray/black). **(B)** The individual points are aggregated to individual regions in brain slices ($n_{slices}=4$) that were then used to look at heterogeneities at the level of biological replicates ($n_{mice,treatment}=4$) for each treatment group compared to the vehicle cohort (n = 3). Horizontal dashed lines reflect the treatment grouped averages to demonstrate a trend that matches the change in spread and enclosed area of the individual cells. **(C)** These values were then aggregated to a per mouse basis and statistical analyses were performed to compare treatment groups to recapitulate temporal trends in IBA1 expression levels observed at the ensembled aggregate level at a per-cell basis.

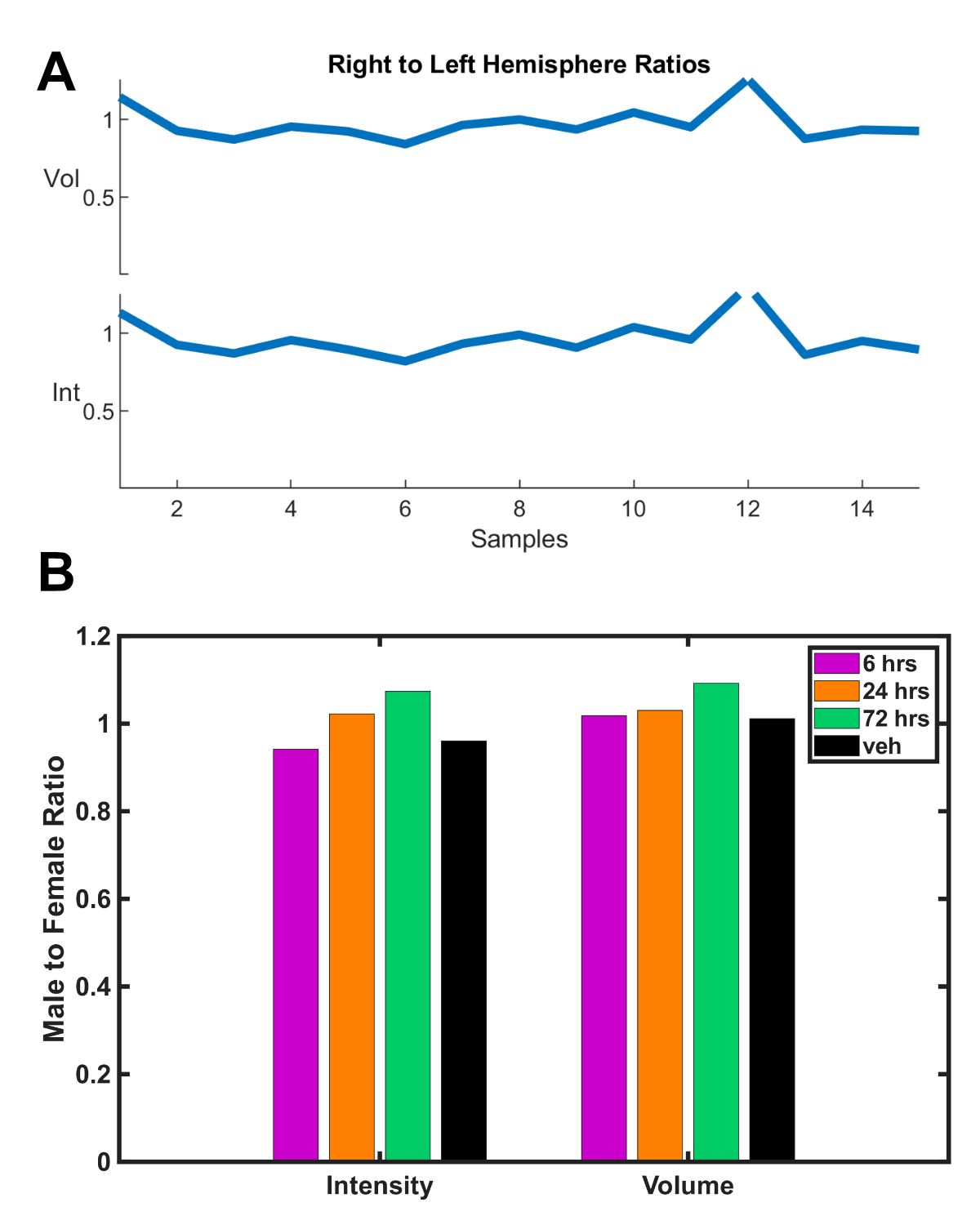

#### Figure S4. Neuroinflammatory activation of microglia does not demonstrate hemispheric preference nor sex-based differences.

**(A)** Comparing measured values from IBA1+ objects in left and right hemispheres to confirm that bilateral administration of i.n. LPS resulted in similar activation profiles in both hemispheres. **(B)** Comparing measured values between biological duplicates based on sex within the same treatment group to confirm no sex-based differences within the scope of the study to allow for aggregating measurements from both sexes.

### TABLES

| Table S1. Selection of unconjugated primary antibodies and dilutions used for immunofluorescent staining | | | | | |
| --- | --- | --- | --- | --- | --- |
| **Sample** | **Host** | **Target** | **Dilution** | **Source** | **Catalog #** |
| IHC  Brain Sections | Rabbit | CD106/ VCAM-1 | 1:200 | MilliporeSigma | CBL1300 |
|  | Rat | Iba-1 | 1:1,000 | Wako | 019-19741 |
|  | Rat | CD68 | 1:1,1000 | BioRad | MCA1957GA |
|  | Chicken | Homer-1 | 1:500 | Synaptic Systems | 160006 |
|  | Mouse | PSD-95 | 1:500 | NeuroMab | 75-028 |
|  | Hamster | CD31/ PECAM-1 | 1:250 | MilliporeSigma | MAB1398Z |
|  | Goat | Fibrinogen | 1:200 | Dako | A0080 |
|  | Rabbit | GLUT-1 | 1:250 | Abcam | 15309 |
| ICC Murine MPS | Rabbit | CD106/ VCAM-1 | 1:250 | MilliporeSigma | CBL1300 |
|  | Biotin | Neutrophil 7/4 | 1:200 | LS Bio | LS-C344780-100 |
|  | Hamster | CD31/ PECAM-1 | 1:250 | MilliporeSigma | MAB1398Z |
|  | Rat | Histone H3 citrunillated | 1:250 | Novus Biologicals | NB100-57135 |

| Table S2. Selection of fluorophore-conjugated antibodies used for immunofluorescent staining | | | | | |
| --- | --- | --- | --- | --- | --- |
| **Host** | **Reactivity** | **Fluor** | **Dilution** | **Source** | **Catalog** |
| Goat | Rabbit | Alexa 568 | 1:750 | ThermoFisher | A-11036 |
|  | Rat | Alexa 647 |  | ThermoFisher | A-21247 |
|  | Mouse | Alexa 488 |  | ThermoFisher | A-11006 |
|  | Chicken | Alexa 647 |  | ThermoFisher | A-21449 |
|  | Rabbit | Alexa 488 |  | ThermoFisher | A-11008 |
|  | Mouse | Alexa 647 |  | ThermoFisher | A-21235 |
| Goat | Hamster | Alexa 488 | 1:500 | JacksonImmuno | 127-545-099 |
| Avidin | Biotin | QDot 605 | 1:500 | ThermoFisher | Q10103MP |
| Rat | Ly6G/Ly6C | Alexa 700 | 1 μL/rxn | ThermoFisher | 56-5931-82 |
| Rat | CD11b | PE-TexasRed |  | ThermoFisher | RM2817 |

| Table S3. List of Reagents, Materials, and Cell Sources | | |
| --- | --- | --- |
| **Item** | **Manufacturer / Supplier** | **Catalog** |
| LPS (E. coli strain O111:B4) | Millipore Sigma | L2630 |
| Mouse Direct PCR Kit | Fisher Scientific | B40015 |
| Protease Inhibitor Cocktail tablets | Roche | 12352200 |
| Lysing Buffer | BD Biosciences | 555899 |
| Diff-Quick | Siemens | 10736133 |
| RNeasy Lipid Tissue Mini Kit | Qiagen | 74804 |
| High-Capacity RNA-to-cDNA Kit | ThermoFisher | 4387406 |
| Bovine Serum Albumin | MilliporeSigma | A8806 |
| Ultrapure EDTA | ThermoFisher | 15575020 |
| Trypan Blue | ThermoFisher | 15250061 |
| Fc-block | ThermoFisher | 14-9161-73 |
| FITC-Dextran 2,000 kDa | ThermoFisher | D7137 |
| FITC-Dextran 40 kDa | ThermoFisher | D1845 |
| Triton X-100 | ThermoFisher | 85111 |
| Prolong Diamond Antifade Mountant | ThermoFisher | P36965 |
| IL-1β | PeproTech | 211-11B |
| ΤΝFα | PeproTech | 315-01A |
| Dulbecco’s Modified Eagle Medium | ThermoFisher | 10567022 |
| Fetal Bovine Serum | Atlas Biologicals | F-0500-D |
| 1-Step Polymorphs | Fisher Scientific | NC9189798 |
| MojoSort Mouse Neutrophil Isolation Kit | BioLegend | 480058 |
| 0.25% Trypsin-EDTA | ThermoFisher | 25200056 |
| Poly-D-Lysine | MilliporeSigma | P1149 |
| Collagen IV | MilliporeSigma | C5533 |
| Fibronectin | MilliporeSigma | F1141 |
| Human endothelial serum-free media | ThermoFisher | 11111-044 |
| B-27 supplement | ThermoFisher | 17504044 |
| Human fibroblast growth factor (h-FGF) | Tocris | 233-FB-500 |
| Glycine | BioRad | 1610718 |
| Tween-20 | MilliporeSigma | P1379 |
| Prolong Diamond Antifade Mountant with DAPI | ThermoFisher | P36962 |
| μSiM Components | SiM Pore | NPSN100-1L |
| Human iPSCs | WiCell | iPS(IMR90)-4 |
| Murine immortalized brain microvascular endothelial cells (bEnd.3) | ATCC | CRL-2299 |

| **Table S4. TaqMan® Gene Expression Assay kits for murine qPCR Analysis** | |
| --- | --- |
| **Target** | **Assay ID** |
| Actb | Mm02619580_g1 |
| Ly6g | Mm04934123_m1 |
| Mpo | Mm01298424_m1 |
| Il1b | Mm00434228_m1 |
| Tnfa | Mm00443258_m1 |
| Ccl2 (Mcp1) | Mm00441242_m1 |
